# Resistance to miltefosine results from amplification of the *RTA3* floppase or inactivation of flippases in *Candida parapsilosis*

**DOI:** 10.1101/2021.12.16.473093

**Authors:** Sean A. Bergin, Fang Zhao, Adam P. Ryan, Carolin A. Müller, Conrad A. Nieduszynski, Bing Zhai, Thierry Rolling, Tobias M. Hohl, Florent Morio, Jillian Scully, Kenneth H. Wolfe, Geraldine Butler

**Affiliations:** School of Biomolecular and Biomedical Science, Conway Institute, University College Dublin, Belfield, Dublin 4, Ireland; Sir William Dunn School of Pathology, University of Oxford, Oxford, UK; Earlham Institute, Norwich Research Park, Norwich NR4 7UZ, UK; Infectious Disease Service, Department of Medicine, Memorial Sloan Kettering Cancer Center, New York, NY, USA; Immunology Program, Sloan Kettering Institute, Memorial Sloan Kettering Cancer Center, New York, NY, USA; Laboratory of Quantitative Engineering Biology, Shenzhen Institute of Synthetic Biology, Shenzhen Institutes of Advanced Technology, Chinese Academy of Sciences, Shenzhen, China; Department of Medicine, Weill Cornell Medical College, New York, NY, USA; Laboratoire de Parasitologie et Mycologie, CHU de Nantes, Nantes, France; Cibles et Médicaments des Infections et du Cancer, IICiMed UPRES EA 1155, UFR de Sciences Pharmaceutiques et Biologiques, 1 rue Gaston Veil, 44035 Nantes, France; School of Medicine, Conway Institute, University College Dublin, Belfield, Dublin 4, Ireland

## Abstract

Flippases and floppases are two classes of proteins that have opposing functions in the maintenance of lipid asymmetry of the plasma membrane. Flippases translocate lipids from the exoplasmic leaflet to the cytosolic leaflet, and floppases act in the opposite direction. Phosphatidylcholine (PC) is a major component of the eukaryotic plasma membrane and is asymmetrically distributed, being more abundant in the exoplasmic leaflet. Here we show that gene amplification of a putative PC floppase or double disruption of two PC flippases in the pathogenic yeast *Candida parapsilosis* results in resistance to miltefosine, an alkylphosphocholine drug that affects PC metabolism that has recently been granted orphan drug designation approval by the US FDA for treatment of invasive candidiasis. We analysed the genomes of 170 *C. parapsilosis* isolates and found that 107 of them have copy number variations (CNVs) at the *RTA3* gene. *RTA3* encodes a putative PC floppase whose deletion is known to increase the inward translocation of PC in *Candida albicans*. *RTA3* copy number ranges from 2 to >40 across the *C. parapsilosis* isolates. Interestingly, 16 distinct CNVs with unique endpoints were identified, and phylogenetic analysis shows that almost all of them have originated only once. We found that increased copy number of *RTA3* correlates with miltefosine resistance. Additionally, we conducted an adaptive laboratory evolution experiment in which two *C. parapsilosis* isolates were cultured in increasing concentrations of miltefosine over 26 days. Two genes, *CPAR2_303950* and *CPAR2_102700*, gained homozygous protein-disrupting mutations in the evolved strains and code for putative PC flippases homologous to *S. cerevisiae DNF1*. Our results indicate that alteration of lipid asymmetry across the plasma membrane is a key mechanism of miltefosine resistance. We also find that *C. parapsilosis* is likely to gain resistance to miltefosine rapidly, because many isolates carry loss-of-function alleles in one of the flippase genes.

**Author summary:** Miltefosine was developed as an anticancer drug but is commonly used to treat infections with the protozoan parasites *Leishmania* and *Trypanosoma cruzi*. More recently, it has been used to treat fungal infections, and in 2021 it was designated as an orphan drug by the US Food and Drug Administration for treatment of invasive candidiasis. Miltefosine is a derivative of phosphatidylcholine (PC), a major constituent of the cell membrane. PC and other phospholipids are asymmetrically distributed across the cell membrane. The mechanism of action of miltefosine is unknown. Here, we show that either increasing the activity of a putative floppase, which controls outward “flop” movement of phospholipids, or decreasing the activity of flippases, which control inward “flip” movement, results in increased resistance of the fungal pathogen *Candida parapsilosis* to miltefosine. This result suggests that miltefosine acts by controlling the localisation of PC or other phospholipids in the membrane. Importantly, we find that many *C. parapsilosis* isolates carry mutations in one flippase gene, which renders them partially resistant to miltefosine, and prone to easily acquiring increased resistance.

## Introduction

Lipids in yeast plasma membranes belong to three main families: sterols, sphingolipids, and phospholipids (or glycerophospholipids) [1]. Phospholipids (including phosphatidylcholine (PC), phosphatidylethanolamine (PE), phosphatidylinositol (PI), and phosphatidylserine (PS)) and sphingolipids (including the long chain bases (LCBs) dihydrosphingosine (DHS) and phytosphingosine (PHS)) are distributed asymmetrically in the plasma membrane bilayer [2, 3]. Plasma asymmetry is determined by the action of lipid translocators via rapid inward (flip) and outward (flop) movements [4]. Most flippases are members of the P4-ATPase family [5], and most floppases are ABC-type transporters [6]. The Rta1 family of 7 transmembrane domain proteins are also predicted lipid transporters [7].

Several antimicrobial drugs are targeted against lipid synthesis, such as azole drugs, which target ergosterol synthesis in fungi [8–10]. Miltefosine (hexadecylphosphocholine) is an alkylphosphocholine derivative that inhibits PC biosynthesis or localization [11]. Miltefosine was initially investigated as an antitumor agent, but it was found to have a very limited anticancer effect [12]. However, it has been used successfully to treat infections with *Leishmania* and *Trypanosoma cruzi* [13–15]. Recently, miltefosine has also been shown to have antifungal activity, including against *Aspergillus* [16], *Cryptococcus* [17] and *Paracoccidioides* [18]. *Anti-Candida* activity has also been reported, including against *Candida auris* [19, 20], *Candida krusei* [21] and *Candida albicans*, where it inhibits growth of both planktonic and biofilm cells [22]. In 2021, miltefosine was granted orphan drug designation approval by the US Food and Drug Administration (FDA) for treatment of invasive candidiasis (https://www.accessdata.fda.gov/scripts/opdlisting/oopd/detailedIndex.cfm?cfgridkey=843921).

The mechanism of action of miltefosine is not completely understood. In liver cells, miltefosine is proposed to reduce the biosynthesis of PC by inhibiting the activity of CTP:phosphocholine cytidylyltransferase (CT) and PE N-methyltransferase activity [23]. Treatment of *Leishmania* with miltefosine decreases the amount of PC and increases PE in the membrane, because of a reduction of intracellular choline [24]. In addition, cytochrome c activity in the mitochondria is inhibited, and the plasma membrane Ca^2+^ channel is disrupted [25]. Studies in *Saccharomyces cerevisiae* suggest that related drugs, such as edelfosine (lysophosphatidylcholine) also disrupt lipid rafts in the cell membrane, perhaps by directly interacting with sterols [26]. In *Aspergillus fumigatus*, miltefosine affects metabolism of lipids and fatty acids, as well as by increasing mitochondrial fragmentation [16].

Most flippases belonging to the P4-ATPase family, which consist of two subunits, a catalytic alpha-subunit with 10 transmembrane alpha-helices, and a beta-subunit with two transmembrane alpha-helices belonging to the Cdc50 family [5]. Flipping of PE, PS and PC is specifically associated with Class 3 P4-ATPases, represented by Dnf1 and Dnf2 in *S. cerevisiae* [27]. Disruption of *DNF1* or *DNF2*, or of *LEM3* (*CDC50*), results in miltefosine resistance in *S. cerevisiae* [28]. Similarly, mutations in P4-ATPases in *Leishmania* species confer resistance to miltefosine [29–32].

In *Candida* species, flippases have not yet been associated with miltefosine resistance. However, deletion of an unrelated gene, *RTA3*, results in increased miltefosine sensitivity in *C. albicans* [33]. Rta3 is a member of the Rta1/Rsb1-like family in *S. cerevisiae*, which encode putative transporters with seven transmembrane domains (TMs). ScRsb1 controls the localization of sphingoid bases in *S. cerevisiae*, including phytosphingosine and dihydrosphingosine [34, 35]. Rsb1 is localized to the membrane and is likely to act as a membrane transporter (a floppase) [7], or possibly as a regulator of transporters [36]. Overexpression of another family member, *ScRTA1*, correlates with increased resistance to 7-amino cholesterol, an inhibitor of 8-7 sterol isomerase and 14-sterol reductase [37]. The Rsb1 family members have not been fully characterised in *Candida* species. As well as showing that deleting *C. albicans RTA3* (*CaRTA3*) increases sensitivity to miltefosine, Srivastava et al [33] also showed that CaRta3 localizes to the plasma membrane, and that a fluorophore-labeled PC accumulates in the inner leaflet of the membrane in an *CaRTA3* deletion. They proposed that CaRta3 is either a floppase, or that it inhibits the activity of a flippase.

Some previous reports suggest that *CaRTA3* is also associated with resistance to azoles. Rogers and Barker [38, 39] found that transcription of *CaRTA3* is up-regulated in fluconazole-resistant isolates from AIDS patents, and Coste et al. [40] showed that Tac1p, a transcription factor regulating expression of the MDR transporters *CDR1* and *CDR2*, also regulates expression of *CaRTA3* via binding to the drug-response element. Whaley et al [41] reported that deleting *CaRTA3* reduced fluconazole resistance of a clinical isolate of *C. albicans*, and that overexpressing *CaRTA3* increased resistance. However, Srivastava et al. [33] showed that deleting *CaRTA3* in a reference isolate had no effect on fluconazole resistance. The role of Rta3 in azole resistance is therefore unclear.

In 2020, we found that the *RTA3* gene has undergone amplification in 23 clinical isolates of *Candida parapsilosis*, resulting in increased copy number of 24-33x [42]. Variable copy numbers of *RTA3* were also found by West et al [43] in four isolates. Here, by genome sequencing, we report that *RTA3* amplification is very common in *C. parapsilosis*. We find 16 distinct amplifications of *RTA3* with unique endpoints, indicative of independent and parallel amplification events. We also find a fusion to a related neighboring gene, *RTA2*. Copy number of *RTA3* is correlated with resistance to miltefosine, but not to azoles. In addition, we find that resistance to miltefosine induced by exposure to the drug is associated with loss-of-function mutations in two P4-ATPase flippase genes.

## Results

### *C. parapsilosis* phylogeny shows independent origins of *RTA3* CNVs

We explored the relationships of 170 *Candida parapsilosis* isolates using a genome-wide SNP alignment approach, generating the largest phylogeny of *C. parapsilosis* isolates to date (Fig 1). Isolates were obtained from several locations, including some previously published, and 127 that are sequenced and described here for the first time (S1 Table). Most isolates originated from the Memorial Sloan Kettering Cancer in New York (designated by MSK) and the Centre Hospitalier Universitaire de Nantes, France (designated by FM) (Fig 1).

**Figure 1.**
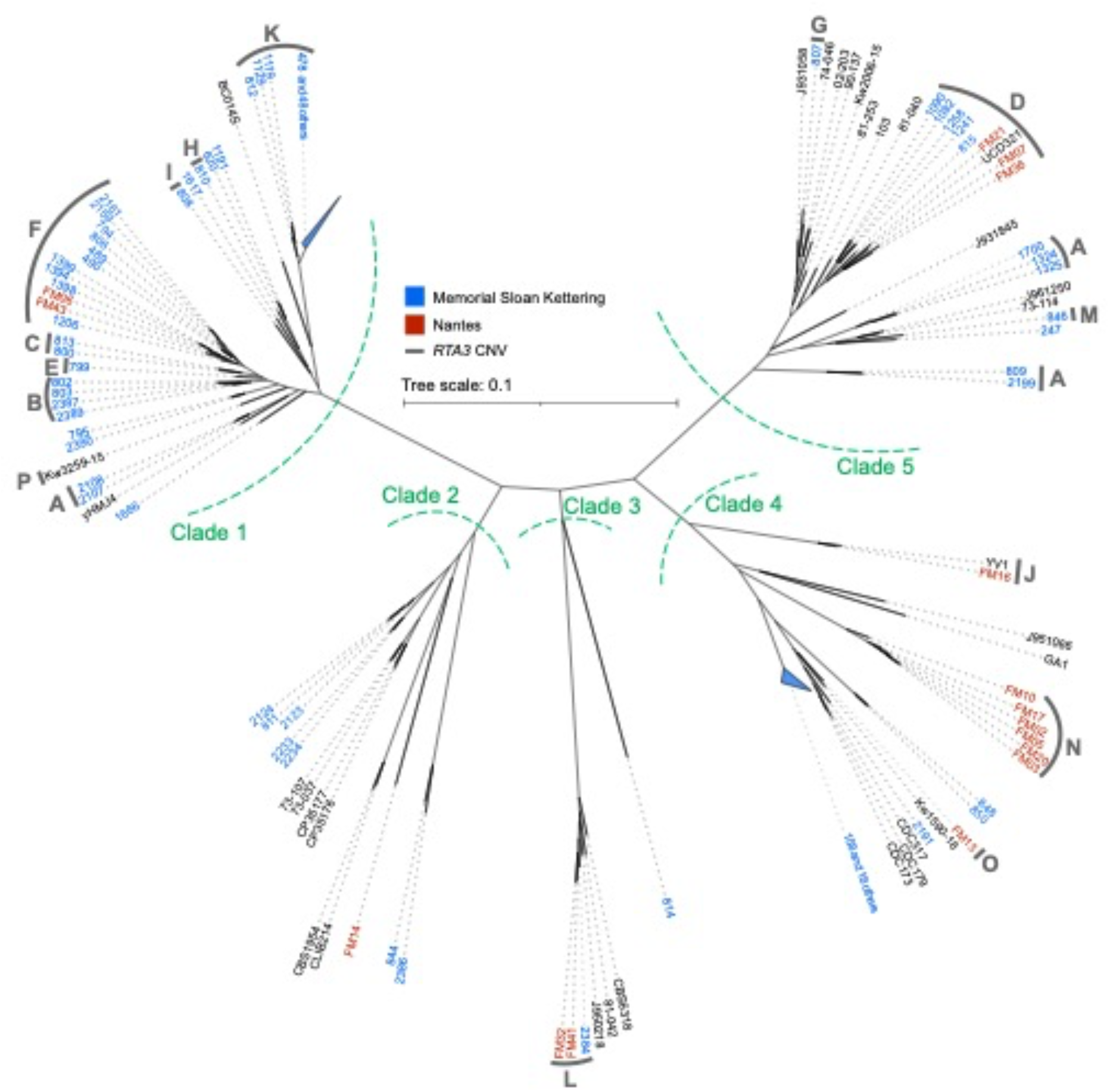
SNP-based unrooted phylogeny of 170 *C. parapsilosis* isolates. Isolates harboring a CNV at *RTA3* are marked with a thick gray bar and a letter (A-P) corresponding to one of the 16 CNVs found. Isolates from Memorial Sloan Kettering Cancer Center (MSK) are named in blue (the prefix MSK is omitted from these strain names), and isolates from CHU de Nantes (FM) are named in red. Green dashed lines label each of five apparent clades. Fifty similar isolates in Clade 1, and 19 similar isolates in Clade 4, were grouped and are shown as blue triangles. The phylogeny was constructed by calling SNPs for each sample using the GATK HaplotypeCaller tool [44]. Filtered heterozygous sites were resolved using 100 iterations of Random Repeated Haplotype Sampling to provide haploid inputs for tree construction [45]. The tree was then constructed using RAxML with the GTRGAMMA model of nucleotide substitution.

Isolates fall into five major clades, with almost half (84/170), belonging to clade I. Most clade I isolates were isolated at MSK. Half (8/16) of the FM isolates are found in Clade 4. There are 49 highly similar isolates in Clade 1, and 19 highly similar isolates in Clade 4, all from patients at MSK. These may have recently originated from two single isolates, and they are indicated with blue triangles in Fig 1. Some other isolates (e.g. MSK2233 and MSK2234 in Clade 2) may also share a recent origin. However, overall there is little evidence for geographical clustering. Each clade includes at least one isolate from both MSK and FM (Fig 1). Three isolates from a clinical setting in Kuwait [46] are each located in a different clade (designated by Kw in Fig 1). Interestingly, an environmental sample isolated from Irish soil, UCD321, groups with clinical isolates from both MSK and FM (in Clade 5). The diversity of isolates obtained from the same clinical setting, and the close relationship between isolates from different geographical settings, highlight the global nature of *C. parapsilosis* as a human pathogen.

We previously showed that the *RTA3* gene, encoding a putative floppase, had undergone extensive copy number amplification in 23 *C. parapsilosis* isolates [42]. This was also observed in a small number of isolates by West et al [43]. We therefore characterized copy number variants (CNVs) at the *RTA3* locus in all sequenced isolates. We identified increased *RTA3* copy numbers in 104 of 170 isolates. Sixteen CNVs with unique endpoints were observed (assigned letters A-P, Figs. 1, 2). Nine different CNVs were observed in isolates from Clade 1, whereas only CNV-L is found in isolates in Clade 3, and there are no CNVs in Clade 2 isolates (Fig 1). Most (15/16) of the CNVs have a single evolutionary origin, and some are present in only a single isolate (Fig 1). However, CNV-A has three separate origins on the tree (one in Clade 1, and two in Clade 5).

**Figure 2.**
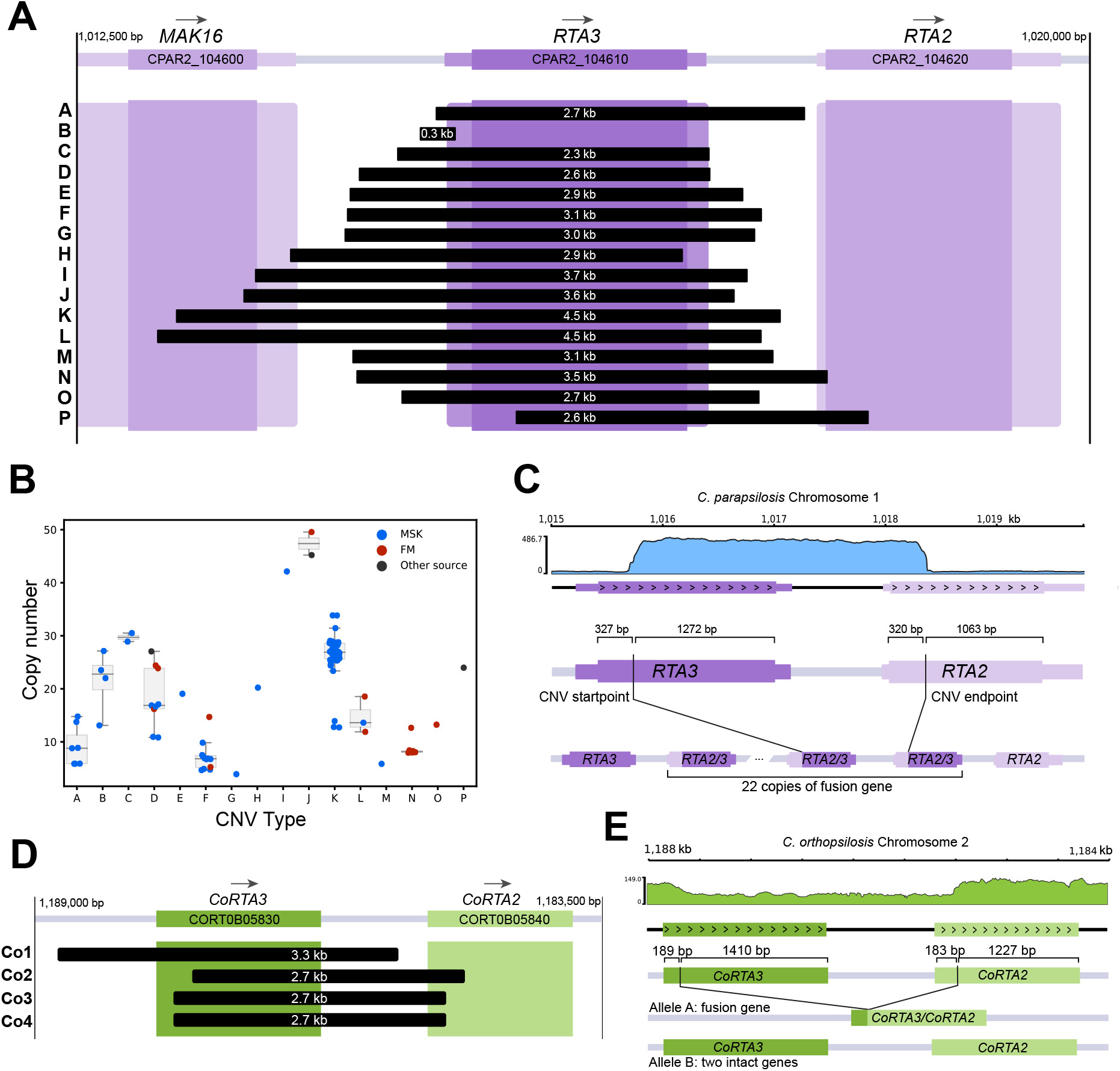
CNVs amplify different sequences at the *RTA3* locus. A. Span of 16 different CNVs at the *RTA3* locus in *C. parapsilosis*. The coding sequence of *RTA3* is shown by a dark purple box, with the flanking UTRs in a lighter color. The neighbouring genes *MAK16* and *RTA2* are also shown. The extents of the tandemly repeating units in each of 16 CNVs are shown by black boxes, labelled (A-P) on the left, and their lengths are indicated in white. Exact breakpoints were identified by interrogating split reads from the Illumina data, and the array structure of CNVs D and K was verified by MinION sequencing of isolates UCD321, MSK478, and MSK812. B. Copy number of *RTA3* in each *C. parapsilosis* isolate. Each dot represents a single isolate: blue isolates from MSK, red from CHU de Nantes, and dark grey from other sources. Median and interquartile ranges are shown for CNVs present in more than one strain. Dots are jittered for clarity. C. *RTA2/3* fusion gene in strain 434 containing CNV-P. A coverage plot created by PyGenomeTracks [47] is shown at the top. The schematic shows the position of CNV breakpoints in relation to the CDS of both genes, and the structure of the inferred array of fusion genes. D. Amplifications of *RTA3* in *C. orthopsilosis*. Orientation of the chromosome has been flipped to highlight the similarities to *RTA3* CNVs in *C. parapsilosis*. E. A deletion in *C. orthopsilosis* generates a *CoRTA3/CoRTA2* fusion. A coverage plot of the locus is shown at the top. The schematic shows CNV breakpoints in both *CoRTA3* and *CoRTA2* and how the resulting fusion gene is formed at one allele, leaving the other allele intact.

### 16 unique CNVs amplify *RTA3*

We found that *RTA3* has been amplified in 16 different types of CNV, each with unique endpoints. Thirteen CNV patterns result from tandem duplications that amplify the entire *RTA3* coding sequence (Fig 2A), and these repeat units range in size from 2.3 to 4.5 kb. Some of these CNVs extend into the upstream neighbouring gene *MAK16*, and one (CNV-N) includes the first 17 bp in the coding sequence of the downstream gene *RTA2*. The copy number of *RTA3* varies both in isolates that share the same CNV, and in isolates with different CNVs (Fig 2B). Isolate MSK807 (CNV-G) has the lowest estimated copy number of *RTA3* among isolates with CNVs, at four copies, whereas isolate FM16 (CNV-J) has the highest copy number, at 50 copies.

For CNV-K, three isolates have roughly half the *RTA3* copy number of other isolates with the same CNV (~13x instead of ~26x; Fig 2B; S1 Table). We considered the possibility that only one allele of *RTA3* was amplified in these isolates (*C. parapsilosis* is diploid), and that both alleles were amplified in other isolates. We explored this issue by using long read sequencing (Oxford Nanopore) of *C. parapsilosis* MSK812 which is one of the CNV-K isolates with fewer copies. We found that it has 8 copies of *RTA3* at one allele and 6 copies at the other (S1A Fig). Similarly, sequencing of isolate MSK478 (which also has CNV-K, with higher copy number of *RTA3*) showed that it has at least 11 copies at both alleles (S1B Fig). Therefore the variation in copy number among isolates with CNV-K is due to expansion or contraction of the array in both alleles, and not due to hemizygosity for the amplification.

Two of the *RTA3* CNVs alter the structure of the protein it encodes. The amplification in CNV-P starts within the *RTA3* open reading frame, and ends within the related adjacent gene *RTA2* (Fig 2A,C). This repeat structure generates an array of in-frame *RTA2/RTA3* gene fusions, with the N-terminus derived from *RTA2* and the C-terminus from *RTA3* (Fig 2C). *RTA3* and *RTA2* are very similar genes, which likely facilitated the fusion event. In CNV-H, the Rta3 protein is slightly truncated because this CNV consists of a tandem duplication with an endpoint 31 bp upstream of the stop codon of *RTA3*, resulting in a protein that is 10 amino acids shorter than the wild type.

CNV-B is substantially different from the others because it is an amplification of only a small region (269 bp) that is upstream and completely outside the *RTA3* coding sequence (Fig 2A). Sequencing read coverage analysis suggests that the repeat has a complex organization in which direct repeats are interspersed with inverted copies of a central segment flanked by inverted-repeats, resulting in an *N*:(2*N*-1):*N* copy number pattern (S2A Fig). This repeat structure was confirmed by MinION sequencing of isolates MSK802 and MSK803. The structure of the CNV-B repeat is similar to the DUP-TRP/INV-DUP structure seen in some human CNVs [48] and is reminiscent of amplifications formed via Origin-Dependent Inverted Repeat Amplification (ODIRA) [49]. The ODIRA model proposes that replication errors at origins that are surrounded by short inverted repeats generate extrachromosomal intermediates, which are then integrated at the original site [49]. This results in complex CNVs with repeat units in head-to-head and tail-to-tail arrangements that are similar to the CNV-B structure. To date, ODIRA has only been observed in *S. cerevisiae*, and it is unclear whether it occurs in other yeasts [49]. If CNV-B were caused by ODIRA, we would anticipate that the central region of the repeat would contain an origin of replication (S2B Fig). However, determined the temporal order of replication in *C. parapsilosis* using Sort-seq (S2C Fig), and we found no evidence that there is an origin near *RTA3* [50].

We also looked for evidence of *RTA3* amplification in other *Candida* species. *RTA3* is not amplified in 200 sequenced *C. albicans* genomes [51, 52]. However, we identified four different *RTA3* CNVs among 36 strains of *Candida orthopsilosis*, a close relative of *C. parapsilosis* (Fig 2D). Only one of these (CNV-Co1) consists of a simple tandem amplification of the entire *CoRTA3* ORF. Two others, CNV-Co2 and CNV-Co3, amplify a fusion of the *CoRTA2* and *CoRTA3* genes, similar to CNV-P in *C. parapsilosis* (Fig 2D). Interestingly, the final *C. orthopsilosis* CNV (CNV-Co4) involves a deletion. In the two strains containing CNV-Co4, the 3’ end of *CoRTA3*, the 5’ end of *CoRTA2*, and the intergenic space between them are deleted at one allele (Fig 2E). This results in a new fusion gene, with the N-terminus derived from *CoRTA3* and the C-terminus derived from *CoRTA2*. Single copies of *CoRTA2* and *CoRTA3* are intact on the other allele (Fig 2E). This is the only example we have seen of an *RTA3* deletion in *C. parapsilosis, C. orthopsilosis* or *C. albicans*.

### Copy number of *RTA3* is correlated with miltefosine, but not fluconazole, resistance

Deletion of *RTA3* in *C. albicans* has been shown to increase susceptibility to miltefosine [33] and possibly to fluconazole [41]. We therefore investigated the effect of copy number on resistance of *C. parapsilosis*. Fig 3A shows that amplification of *RTA3* is associated with miltefosine resistance. For example, *C. parapsilosis* strains with only two copies of *RTA3* (one at each allele, e.g. the reference strain CLIB214) fail to grow at miltefosine concentrations of 10 μg/ml, whereas all the strains with *RTA3* amplifications can grow at this concentration. Strains with CNVs A, H, I, J and L can tolerate miltefosine concentrations up to at least 30 μg/ml, as can isolates with CNV-B (which amplifies only the region upstream of *RTA3*). The CNV with the weakest effect on MF resistance is CNV-G, which can tolerate 10-15 μg/ml, which is still higher than the reference strain CLIB214.

**Figure 3.**
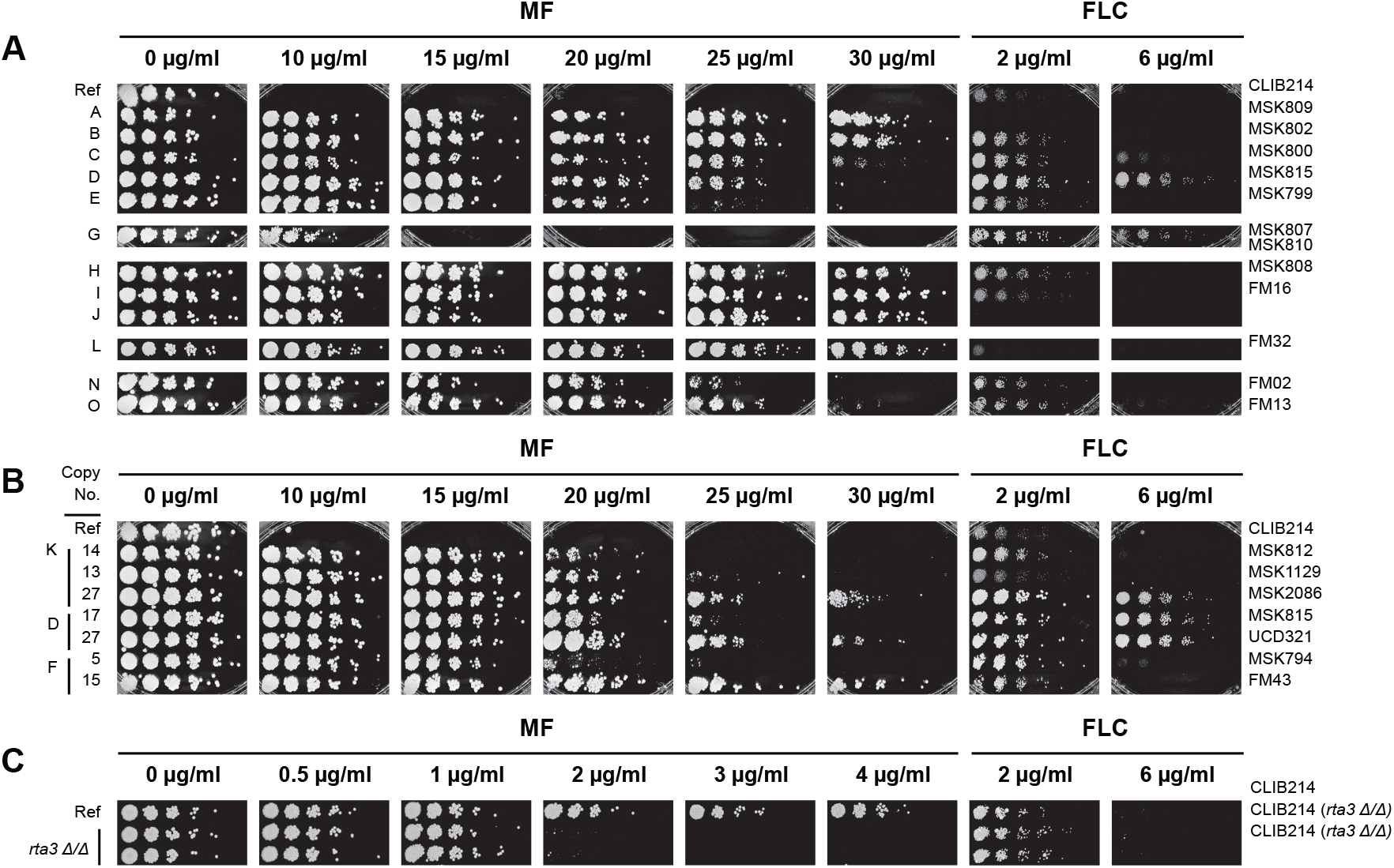
CNVs of *RTA3* correlate with resistance to miltefosine. A. A representative isolate of most CNV patterns (Fig 2) was grown on YPD with increasing concentrations of miltefosine (MF) or fluconazole (FLC) as shown. Cells were serially diluted (1/5). The CNV pattern is indicated on the left, with the strain name shown on the right. MF and FLC plates were incubated for 48 hr. “Ref” indicates the reference strain *C. parapsilosis* CLIB214. B. Copy number is directly correlated with miltefosine resistance. Isolates with different copy numbers of CNV-K, CNV-D and CNV-F were grown as in (A). The *RTA3* copy number of each isolate is shown on the left. C. Deleting *RTA3* reduces resistance to miltefosine. *RTA3* was deleted in *C. parapsilosis* CLIB214 using CRISPR-Cas9. The growth of two biological replicates is shown, as in (A), except that incubations on FLC were for 72 h.

When different isolates carrying the same CNV are compared, miltefosine resistance is seen to be correlated with the copy number of *RTA3* (Fig 3B). For example, *C. parapsilosis* isolates MSK812 and MSK1129, which have 13-14 copies of CNV-K, can tolerate miltefosine concentration up to ~20 μg/ml, whereas *C. parapsilosis* MSK2086 which has 27 copies survives up to 30 μg/ml. Similarly, *C. parapsilosis* MSK815 with 17 copies of CNV-D and MSK794 with 5 copies of CNV-F tolerate miltefosine concentrations up to ~20-25 μg/ml whereas UCD321 and FM43 with 27 and 15 copies of CNV-D and CNV-F respectively survive up to ~30 μg/ml (Fig 3B).

The isolates have variable levels of sensitivity to fluconazole (Fig 3), but the copy number of *RTA3* does not correlate with susceptibility to this drug. For example, isolates MSK815 and UCD321 which have 17 and 27 copies of CNV-D differ in their susceptibility to miltefosine, but they both tolerate fluconazole levels of 6 μg/ml.

We also found that deleting *RTA3* in the reference isolate of *C. parapsilosis* (CLIB214) by CRISPR-Cas9 editing [53] results in increased sensitivity to miltefosine (Fig 3C). Very low levels of miltefosine were used (up to 4 μg/ml), because the parental strain, without any *RTA3* amplifications, is very sensitive. However, susceptibility to fluconazole was not affected by deleting *RTA3* (Fig 3C).

Increasing the *RTA3* copy number correlates with increased expression, as shown by West et al [43]. However we wondered whether amplifying the upstream region (CNV-B) has the same effect as amplifying the entire open reading frame. To explore this further, we measured *RTA3* expression by RT-PCR in one isolate of *C. parapsilosis*, MSK808, with approximately 42 copies of *RTA3* (CNV-I), and two isolates (MSK802 and MSK803) in which only the promoter region is amplified (CNV-B). We found that *RTA3* expression is approximately 22-fold higher in *C. parapsilosis* MSK808, and 2.8- to 6.6-fold higher in *C. parapsilosis* MSK802 and MSK803 than in the reference strain CLIB214 (Table 1). There are 28 copies of the direct repeat in CNV-B in *C. parapsilosis* MSK802 and 24 in *C. parapsilosis* MSK803. Thus, *RTA3* promoter region amplification (CNV-B) can lead to increased expression.

**Table 1.** Expression of *RTA3* in strains with different amplifications

| Isolate | Relative Expression | Range | P Value |
| --- | --- | --- | --- |
| MSK802 (CNV-B) | 6.57 | 2.12 - 20.35 | 7.37E-03 |
| MSK803 (CNV-B) | 2.79 | 1.24 - 6.25 | 3.39E-02 |
| MSK808 (CNV-I) | 22.50 | 6.52 - 77.62 | 6.92E-05 |
| CLIB214 (Reference) | 1.00 | 0.19 - 5.27 | N/A |

### Generation of miltefosine-resistant strains by experimental evolution

The prevalence of *RTA3* amplification in *C. parapsilosis* isolates suggests that there may be some strong selective pressure inducing amplification. To determine if miltefosine is the driving force, we characterized the effect of exposing isolates to increasing concentrations of miltefosine, in an adaptive laboratory evolution approach. We started with two isolates with only one copy of *RTA3* at each allele that were in the same clade as other isolates that had undergone amplification - *C. parapsilosis* MSK795 from Clade 1, which is related to isolates that have undergone 4 different CNVs (B, C, E, and F), and *C. parapsilosis* MSK247 from Clade 5, which is related to isolates with CNVs A, D, J and M (Fig 1; Fig 4). Five lineages were evolved from MSK247 (247A to 247E) and one from MSK795 (795B). Isolates were cultured in YPD with increasing concentrations of miltefosine, up to a maximum of 32 μg/ml, over a 26-day period (Fig 4A). Sixteen randomly picked evolved colonies from each lineage tolerated miltefosine levels of 40 μg/ml (Fig 4B). The genomes of ten isolates derived from MSK247 and two from MSK795 were sequenced, together with the parental strains. The sequenced strains are listed in Methods.

**Figure 4.**
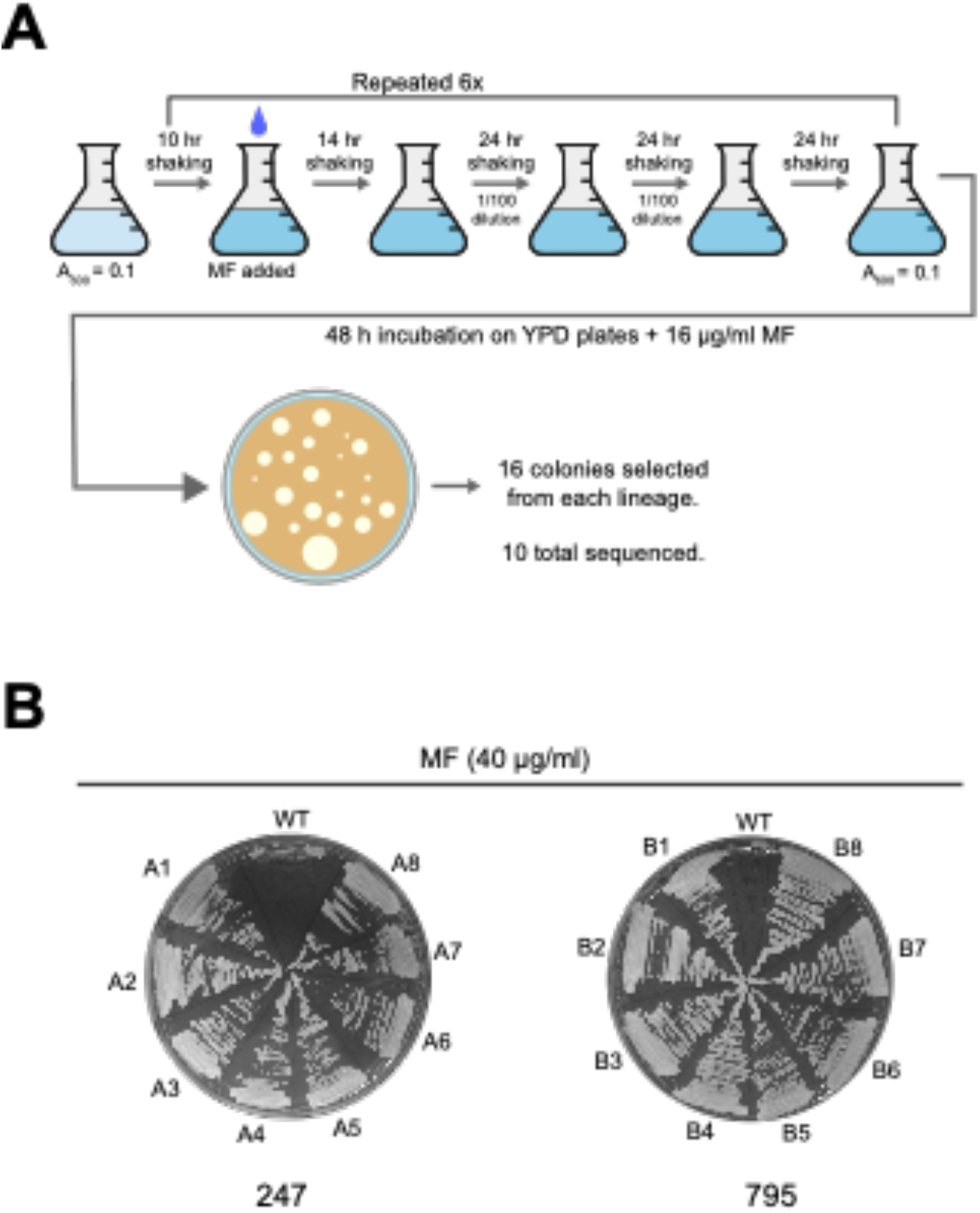
Generation of miltefosine-resistant *C. parapsilosis* isolates by adaptive laboratory evolution. A. Six colonies (5 derived from *C. parapsilosis* MSK247 and one from *C. parapsilosis* MSK795) were inoculated in YPD at A_600_=0.1,and incubated at 30°C for 10 h. Miltefosine (indicated with a blue drop) was added to a final concentration of 1 μg/ml, the cultures were incubated for a further 14 h, and then re-inoculated into the same media using a 1/100 dilution and incubated for 24 h. The dilution was repeated every 24 h for 3 days. Cells were then inoculated in fresh media, grown for 10 h, and the miltefosine concentration was doubled. The process was repeated until the concentration of miltefosine reached 32 μg/ml (24 days). On day 25, the overnight cultures were plated on YPD with 16 μg/ml miltefosine. Sixteen colonies picked from each lineage for further analysis, and the genomes of 10 were sequenced. B. Growth of representative isolates evolved from *C. parapsilosis* MSK247 and *C. parapsilosis* MSK795 on 40 μg/ml miltefosine. WT = parental strain.

### Miltefosine-resistant isolates acquired homozygous disruptions in two flippase genes

We did not find any evidence of amplification of the *RTA3* locus in any of the evolved miltefosine-resistant strains. However, by comparing the sequences of the evolved strains to those of the parental strains, we identified two genes with homozygous loss-of-function variants in all 10 resistant isolates: *CPAR2_102700* and *CPAR2_303950*. These variants include frameshifts, nonsense mutations, and missense mutations that are predicted to be deleterious (Fig 5A). *CPAR2_102700* and *CPAR2_303950* encode putative Class 3 P4-ATPases and are homologs of the PC flippase genes *DNF1* and *DNF2* in *S. cerevisiae* (Fig 5B).

**Figure 5.**
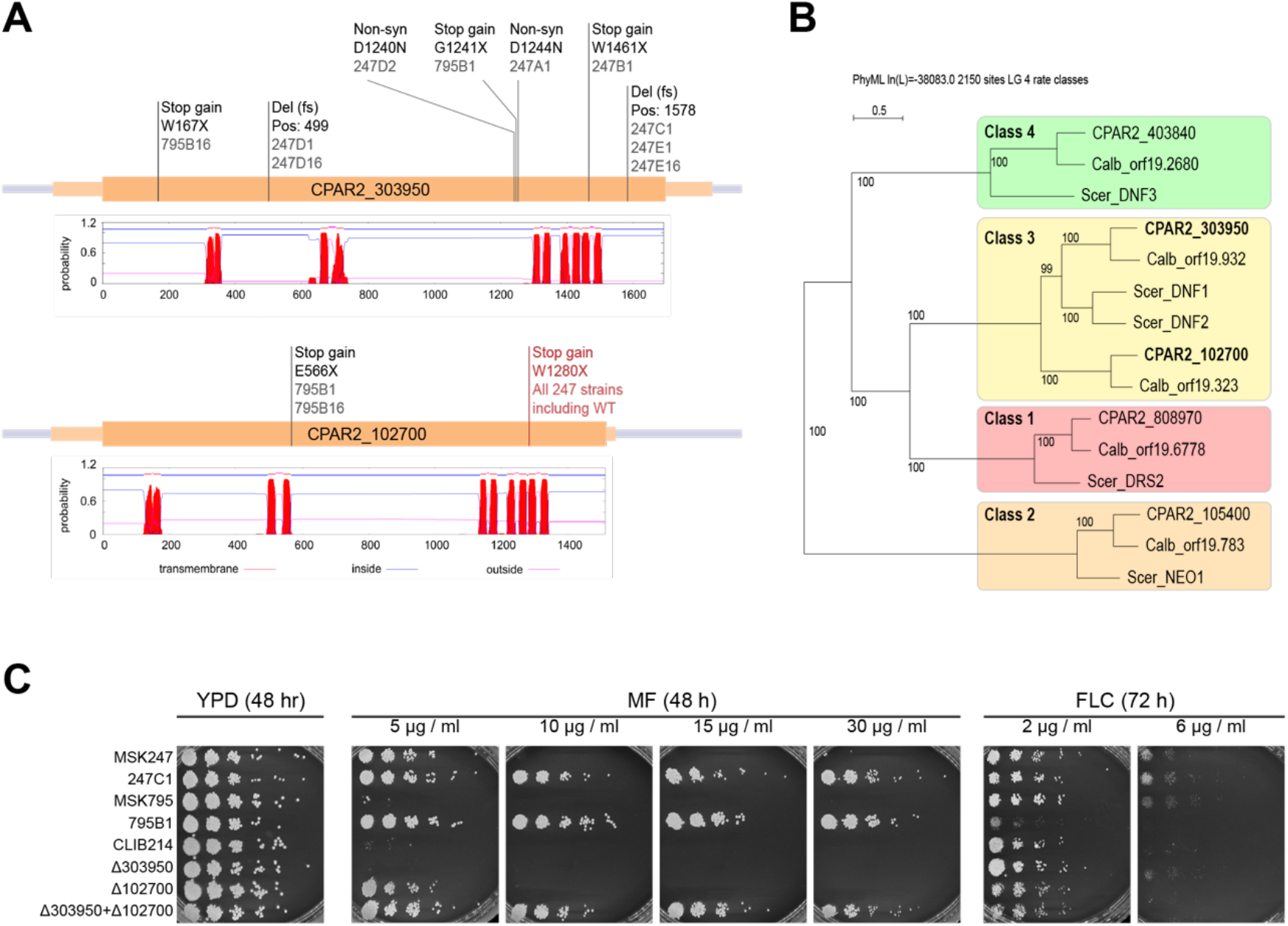
Protein-disrupting mutations in two flippase genes confer resistance to miltefosine in laboratory evolved strains. A. Schematic showing homozygous mutations in P4-ATPases *CPAR2_102700* and *CPAR2_303950*. Mutations named in black were acquired during the laboratory evolution experiment, in the evolved strains named below the mutation name. The mutation named in red is a natural homozygous variant found in the *C. parapsilosis* MSK247 isolate and all strains derived from it. The predicted transmembrane topology of each protein (from TMHMM [54]) is shown, where red peaks show predicted transmembrane domains. B. Phylogeny of P4-ATPases in *C. parapsilosis, C. albicans*, and *S. cerevisiae*. The P4-ATPase genes in three yeast species were identified by BLAST, aligned and a tree was constructed using SeaView [55]. Monophyletic groups of genes were assigned to established classes (1-4) [56]. Both *CPAR2_102700* and *CPAR2_303950* (in bold) belong to Class 3 P4-ATPases. C. Deleting *CPAR2_102700* and *CPAR2_303950* increases resistance to miltefosine, but not to fluconazole. *CPAR2_102700* and *CPAR2_303950* were deleted singly or together in the *C. parapsilosis* CLIB214 background, and growth was compared to isolates evolved from *C. parapsilosis* MSK247 (247C1) and *C. parapsilosis* MSK795 (795B1), as in Figure 3.

The two sequenced isolates derived from *C. parapsilosis* MSK795 (795B1 and 795B16) acquired mutations in both *CPAR2_102700* and *CPAR2_303950*, whereas lineages derived from *C. parapsilosis* MSK247 acquired mutations only in *CPAR2_303950* (Fig 5A). However, subsequent analysis revealed that *C. parapsilosis* MSK247 contains a homozygous natural variant in *CPAR2_102700*, resulting in a Trp-to-Stop nonsense mutation (W1280X; Fig 5A).

*C. parapsilosis* MSK247 tolerates miltefosine concentrations of 5 μg/ml, whereas *C. parapsilosis* MSK795 fails to grow (Fig 5C). Derivatives of both parents that carry homozygous inactivating mutations in both *CPAR2_30395* and *CPAR2_102700* can grow up to 30 μg/ml.

This result suggests that maximum levels of miltefosine resistance requires inactivation of both genes, *CPAR2_30395* and *CPAR2_102700*. To test this hypothesis, we deleted them, both separately and together, in the *C. parapsilosis* CLIB214 genetic background using CRISPR-Cas9 editing (Fig 5C). Deleting *CPAR2_102700* alone in *C. parapsilosis* CLIB214 allows growth up to 5 μg/ml miltefosine, whereas deleting *CPAR2_303950* alone has no effect (Fig 5C). However, strains in which both *CPAR2_303950* and *CPAR2_102700* are deleted tolerate at least 30 μg/ml miltefosine, a similar concentration to strains derived from the experimentally evolved isolates (MSK795B1, MSK247C1) (Fig 5C). Deleting *CPAR2_102700* alone, or in combination with *CPAR2_303950*, slightly increases sensitivity to fluconazole (Fig 5C).

## Discussion

There is some debate over whether Rta3-like proteins are transporters or signaling receptors. Rta3 is a member of the Rta1-family, which encodes proteins with 7 transmembrane domains, similar to the structure of G-protein-coupled receptors (GPCRs) [7]. Within this family, Rta3 is more closely related to *S. cerevisiae* Rsb1 than to other members. Most of the early evidence suggested that Rsb1 directly flips LCBs in the plasma membrane [35]. However, Johnson et al [36] suggested that rather than acting as a transporter, Rsb1 determines resistance to the LCB phytosphingosine by regulating endocytosis of the tryptophan transporter Tat2, either by signalling through a G-protein (as a GPCR), or by an arrestin-mediated effect on targeting of ubiquitin ligases. In addition, Srivistava et al (2017) found that deleting *RTA3* in *C. albicans* resulted in increased flipping of a fluorescently labelled PC, possibly by controlling activity of an unknown flippase [33].

However, several pieces of evidence suggest that Rsb1 is a direct lipid transporter, specifically a floppase. Firstly, Rsb1 shares no sequence similarity with other yeast GPCRs [7]. Secondly, Makuta et al [57] showed that Rsb1 (and not other Rta1-family members) regulates LCB transport, and that this activity is dependent on a loop region following TMS5, and not the C-terminus as expected for a GPCR. Rsb1 in *S. cerevisiae* and Rta3 in *C. albicans* are located in the plasma membrane [33, 34, 41]. The increased flipping observed by Srivistava et al (2017) [33] may be due to crosstalk between sphingolipids and glycerophospholipids (e.g. PC), as shown in *S. cerevisiae* by Kihara and Igarashi (2004) [35]. Changes in the distribution of one may be compensated by changing the distribution of the other [7]. The association of *RTA3* copy number amplification with miltefosine resistance in *C. parapsilosis* is also consistent with a role as a transporter. It is therefore likely that Rta3 is a transporter of LCBs, similar to Rsb1, and that its overexpression and deletion cause changes in the localization of both PC and LCBs. Our results also suggest that *C. parapsilosis CPAR2_102700* and *CPAR2_303950*, members of the class 3 family of P4-ATPases, are the direct flippases of PC. Our data however cannot rule out the alternative model of Johnson et al [36] whereby Rta3 regulates the function of *CPAR2_102700* and *CPAR2_303950*, rather than directly acting as a transporter.

*RTA3* is adjacent to a paralogous gene (called *RTA2*) in many species in the *Candida* (CUG-Ser1) clade [58, 59]. In fact, *C. parapsilosis* CNV-P results from amplification of an *RTA3/RTA2* fusion, with the N-terminus derived from Rta2 and the C-terminus from Rta3 (Fig 2C). Deleting *RTA2* in *C. albicans* increases sensitivity to azole drugs, and overexpression increases resistance [60]. Whaley et al [41] showed that deleting *RTA3* increases sensitivity of an azole-resistant isolate of *C. albicans*, but does not have any effect on a reference isolate. Our results show that *C. parapsilosis RTA3* is unlikely to play any role in azole resistance: deleting the gene does not increase sensitivity, and levels of resistance are not associated with *RTA3* copy number in several different genetic backgrounds. However, the sequence similarity between *RTA2* and *RTA3* in *C. parapsilosis* (and in the related species *C. orthopsilosis* and *C. metapsilosis*) is much higher than between *C. albicans RTA2* and *RTA3* (S3A Fig). This similarity, and the phylogenetic relationship among these sequences (S3A Fig), indicates that in the ancestor of *C. parapsilosis/C. orthopsilosis/C. metapsilosis*, a gene conversion event occurred in which *RTA3* overwrote *RTA2*. The biggest difference between Rta3 and Rta2 in *C. parapsilosis* is that the former has a longer C-terminus (S3B Fig). Because we have never observed amplifications of *RTA2* in miltefosine-resistant *C. parapsilosis* isolates, we suggest that the two genes may have different functions, and that the difference may be related to the C terminus. In *C. albicans*, *RTA2* (and possibly *RTA3*) also contribute to tunicamycin-induced ER stress, which has not been tested in *C. parapsilosis* [61, 62].

Of the 14 CNVs resulting in tandem array amplification of *RTA3*, only CNV-A and CNV-E have obvious repeat sequences at the CNV endpoints, direct repeats of 19 bp and 10 bp respectively. The other 12 either have no repeats, or the repeat is <4 bp. The longer repeat sequence at the breakpoints of CNV-A may explain why it has originated independently three times (Fig 1). CNV-A and CNV-E may result from non-allelic homologous recombination between the direct repeats [48]. It is likely that CNV-P also originated by non-allelic homologous recombination due to the high sequence similarity between *RTA3* and *RTA2*. CNVs with shorter direct repeats at their endpoints may result from microhomology-mediated break induced replication [63]. However, there is no obvious mechanism for CNVs with no repeats at the breakpoints.

CNV-B (upstream from the coding sequence) is particularly intriguing, because it results in inversions and triplications of part of the repeated sequence (S2A Fig). This structure is reminiscent of the origin-dependent ODIRA model for CNV generation [49]. We did not find any evidence of a replication origin in the amplified sequence (S2A Fig), but it remains possible that a rarely used origin is present. It is likely that the amplified region contains a promoter because expression of *RTA3* is increased (Table 1).

Our results strongly suggest that there is selective pressure driving amplification of *RTA3* in *C. parapsilosis*. This conclusion is based on our observation that amplification occurred in isolates from diverse genetic and environmental backgrounds, distributed across the *C. parapsilosis* phylogeny (Fig 1). We also found that there have been multiple independent events of amplification, with at least 16 unique endpoints. West et al (2021) previously described *RTA3* amplifications in 4 *C. parapsilosis* isolates, including one from the New York subway [43]. Those amplifications also have different endpoints, but because the data comes from metagenomics analyses, we cannot determine if their endpoints differ from the 16 CNVs we describe.

The CNVs sometimes encompass parts of the adjacent genes, but *RTA3* is the only complete gene amplified in all of them (except for CNV-B, where its likely promoter is amplified, and CNV-H where its 3’ end is slightly truncated in the amplified copies). The majority of the isolates in this study originated from clinical sources, although one with an *RTA3* amplification (*C. parapsilosis* UCD321) has an environmental origin (it was isolated from soil). There are several known examples of environmental conditions that induce or selecting for CNVs, such as copper resistance resulting from amplification of *CUP1* in *S. cerevisiae* [64, 65], and CNVs associated with growth of *C. albicans* in the presence of fluconazole [66]. It is highly unlikely that the *RTA3* amplifications we discovered were driven by exposure to miltefosine because it is not a commonly used antifungal drug and, as we have shown, exposure to miltefosine is more likely to result in loss-of-function mutations in flippase genes. At present, we do not know what kind of selective pressure led to the widespread amplifications of *RTA3* in *C. parapsilosis*.

Miltefosine has potential as an antifungal drug [20–22], and it has recently been designated as an orphan drug for the treatment of invasive candidiasis (https://www.accessdata.fda.gov/scripts/opdlisting/oopd/detailedIndex.cfm?cfgridkey=843921). However, our analyses suggest that it would be a poor choice, at least for invasive *Candida parapsilosis* infections. Many isolates are naturally resistant because of amplifications of *RTA3*, and in others, resistance rapidly arises due to loss-of-function mutations in the flippase genes. Although mutations in two flippase genes are required for resistance, we note that many of the *C. parapsilosis* samples we examined (56 of 170) have predicted loss-of-function variants in *CPAR2_102700*, meaning that acquiring a mutation in *CPAR2_303950* would be sufficient to render them highly resistant to miltefosine.

## Materials and methods

### Strains and growth

Isolates used are listed in S1 Table. Isolates were maintained on YPD agar (1% Bacto Yeast Extract (212750, Sigma), 2% Bacto Peptone (211677, Sigma), 2% Bacto Agar (214010, Sigma), 2% D-(+)-Glucose (G8270, Sigma)), and liquid cultures were grown in 5 ml YPD broth without agar at 30°C and 200 rpm shaking overnight. For serial dilutions, 0.5 ml of the overnight cultures were harvested at 13,000 rpm at room temperature for 1 min, washed twice in 0.5 ml of phosphate-buffered saline (PBS) buffer (BR0014G, Thermofisher), resuspended in 0.5 ml PBS, and diluted to A_600_= 0.0625 (approximately 6.25 × 10^5^ cells/ml) in PBS. Five-fold serial dilutions were made in PBS and transferred with a pinner to YPD agar plates containing miltefosine (M5571, Sigma Aldrich) or fluconazole (F8929, Sigma Aldrich) at the indicated concentrations. Pinned plates were incubated at 30°C for the indicated times and photographed using a Singer PhenoBooth.

### Gene deletions/disruptions

The entire open reading frames of *CPAR2_104610* (*RTA3*) and the flippases *CPAR2_303950* and *CPAR2_102700* were deleted using CRISPR-Cas9 with the pCP-tRNA system, as described in Lombardi et al [53]. All primers used for gRNA and repair template synthesis are listed in S2 Table.

### Microevolution of miltefosine resistant isolates

The microevolution method was modified from Papp et al [67] and Ene et al. [68]. Three colonies from *C. parapsilosis* MSK247 and *C. parapsilosis* MSK795 were originally chosen. Subsequent analysis showed that 5 evolved lineages were derived from *C. parapsilosis* MSK247 (247A to E), and one from *C. parapsilosis* MSK795 (795B). Colonies were incubated for 10 h in 5 ml YPD at 200 rpm and 30°C. Miltefosine was added to a final concentration of 1.0 μg/ml, and the cultures were incubated for a further 14 h. The cultures were diluted (1:100) into fresh media containing the same concentration of miltefosine every 24 h for 3 days. Overnight cultures were then diluted to an A_600_ = 0.1, incubated for 10 h, and the miltefosine concentration was doubled. The 5-day cycle was repeated until a concentration of 32 μg/ml of miltefosine was reached (see Fig 4A). Cultures were diluted and plated on YPD agar plates containing 16 μg/ml miltefosine, yielding 100-200 colonies each plate, and were incubated at 30°C for 48 h. The genomes of ten resistant isolates (247A1, 247B1, 247C1, 247D1, 247D2, 247D16, 247E1, 247E16, 797B1 and 795B16), together with the parental strains were sequenced by Beijing Genomic Institute via DNBseq.

### Illumina sequencing

Illumina sequencing of MSK isolates was carried out as described in Zhai et al [42]. 45 *C. parapsilosis* isolates from Centre Hospitalier Universitaire de Nantes, France were screened for resistance to miltefosine, and 16 isolates were chosen for sequencing. Genomic DNA was isolated from 5 ml overnight cultures in YPD at 30°C using phenol-chloroform-isoamyl alcohol extraction. Cell pellets were resuspended in 200 μl of extraction buffer (Triton X 100 2% m/v, NaCl 100mM, Tris 10mM pH 7.4, EDTA 1mM, SDS 1% m/v), transferred to screw cap tubes, and approximately 0.3 g of acid-washed beads and 200 μl of Phenol-chloroform-isoamyl alcohol (25:24:1) was added. Cells were lysed using a 1600 MiniG bead beater from Spex SamplePrep for 6 x 30 seconds with 30 s pauses in between, and then centrifuged at 14000 rpm for 10 min at room temperature. The supernatant was extracted twice more with 200 μl of TE and 200 μl of phenol-chloroform-isoamyl alcohol and one 30 s agitation in the bead beater. DNA was precipitated using 80 μl of ammonium acetate (7.5M) and 1 ml of 100% isopropanol, washed with 1 ml of 70% ethanol and air-dried. Pellets were resuspended in 400 μl of TE with 1 μl of RNAse A (100 mg/ml) incubated overnight at 37°C, DNA was precipitated again, and resuspended in 150 μl water. Illumina sequencing was performed by the UCD Conway Genomics Core using a NextSeq 500. 1 ng of gDNA was tagmented (fragmented and tagged with adapter sequences) using the Nextera kit transposome, Dual-indexed paired-end libraries were prepared using Nextera XT DNA Library Prep kit. An Illumina NextSeq500 mid output 300 cycle sequencing kit was used to prepare and run the flowcell (HVGWJAFX2).

### Oxford Nanopore sequencing

Strains were grown overnight in 50 ml YPD broth and genomic DNA was extracted from approximately 4×10^9^ cells using a QIAGEN Genomic-tip 100/G kit (10223, QIAGEN) with minor modifications. The lyticase incubation was extended to 2 h, and the proteinase K incubation was extended to overnight (~15 h). DNA libraries were prepared using three different kits as per manufacturer’s instructions. Libraries from *C. parapsilosis* MSK812 and UCD321 were prepared using a Ligation Sequencing Kit (SQK-LSK109, Oxford Nanopore) using 1 μg of DNA per strain. DNA was repaired using a NEBNext FFPE DNA Repair Mix (M6630, New England Biolabs) and NEBNext Ultra II End repair/dA-tailing Module (E75460, New England Biolabs). Adapters were ligated using NEBNext Quick Ligation Module (E60560, New England Biolabs). Libraries were sequenced on a MinION Mk1C device. Priming, loading and washing was carried using the EXP-FLP002, and SQK-LSK109 and EXP-WSH002 Oxford Nanopore kits respectively as per manufacturer’s instructions. The genomes were sequenced in parallel on one flow cell; the first isolate (*C. parapsilosis* MSK812) was sequenced for 24 h, and the flow cell was washed, re-primed and loaded with the *C. parapsilosis* UCD321 isolate for an additional ~48 h. Libraries from *C. parapsilosis* MSK802 and MSK803 were prepared using a Rapid Barcode Sequencing Kit (SQK-RBK004, Oxford Nanopore) using 400 ng of DNA per sample. Samples were multiplexed using barcodes R04 and R05 respectively. Libraries were sequenced on an original MinION device for 72 h. Priming and loading was performed using the EXP-FLP002 and SQK-RBK004 kits respectively. The *C. parapsilosis* MSK478 library was prepared using a Rapid Sequencing Kit (SQK-RAD004, Oxford Nanopore) using 400 ng of DNA, and sequenced for 72 h using an original MinION device. Basecalling was performed for all samples using guppy_basecaller with the following parameters: “-- input_path fast5 --save_path fastq -c dna_r9.4.1_450bps_fast.cfg --verbose_logs -- cpu_threads_per_caller 5 --num_callers 10”. Guppy v.4.2.2+effbaf8 was used for samples MSK812 and UCD321 and Guppy v.3.6 was used for samples MSK802 and MSK803. For multiplexed samples MSK802 and MSK803, demultiplexing was performed using guppy_barcoder with the following parameters: “--barcode_kits SQK-RBK004 -t 30 -- verbose_logs --trim_barcodes”. Reads were filtered using NanoFilt with the following parameters: “-l 1000 -q 7” [69].

### Sequence analysis

Illumina reads were trimmed with Skewer version 0.2.2 using tags “-m pe -t 4 -l 35 -q 30 -Q 30” [70]. The trimmed reads were aligned to the *C. parapsilosis* reference genome using bwa mem version 0.7.12. The resulting BAM files were sorted and duplicate reads were marked using GenomeAnalysisToolkit (GATK version 4.0.1.2) SortSam and MarkDuplicates tools respectively [44]. Variants were called with GATK HaplotypeCaller using the tag “--genotyping_mode DISCOVERY”, combined using GATK CombineGVCFs and joint-genotyped using GATK GenotypeGVCFs. Variant files were filtered for read depth (<15) using and genotype quality (<40) using GATK VariantFiltration. Additionally, clusters of SNPs (5 SNPs in a 100bp window) were filtered using GATK VariantFiltration. A custom script was used to remove variants that were flanked on either side by a long string of mono- or dinucleotide repeats, and variants that were called as heterozygous but had an allele depth ratio <0.25 or >0.75 (https://github.com/CMOTsean/milt_variant_filtration). Additionally, for tree construction, indels were excluded using GATK SelectVariants with the tag “--select-type-to-include SNP”. For analysis of the evolved strains, a custom script was used to filter out variants in the evolved strains that were also present in the respective parent strain (https://github.com/CMOTsean/milt_variant_filtration).

### SIFT4G analysis

A SIFT prediction database was created for *C. parapsilosis* using the SIFT4G algorithm [71], and the recommended Uniref90 database as a reference for protein sequences [72]. The *C. parapsilosis* prediction database was used to annotate variants from the evolved strains with whether they are likely to be deleterious to protein function.

For each annotated gene in *C. parapsilosis*, the number of evolved strains which carried a variant predicted to be protein function-affecting by SIFT in that gene was tallied. Variants were also visualised using Integrative Genomics Viewer (IGV) to manually check results [73].

### Phylogeny construction

Called SNPS were concatenated and heterozygous sites were resolved randomly to either allele by 1000 iterations of Random Repeated Haplotype Sampling [45]. SNP trees were then constructed from each of the 1000 haploid inputs, using RAxML (v8.2.12) with the GTRGAMMA model of nucleotide substitution and the random number seed “-p 12345” [74]. The tree with the highest maximum likelihood score was chosen, and the remaining 999 trees were used to generate branch support values.

### Estimating copy number

For each strain with a CNV at *RTA3*, the average coverage across the *RTA3* ORF was found using BEDTools coverage (v2.29.2) [75]. This value was divided by the average genome coverage (found with BEDTools genomecov) and multiplied by two to adjust for ploidy, to calculate an estimate for the copy number of *RTA3*. For CNVs that did not cover the entire *RTA3* ORF, the average coverage of a representative section of the CNV was used instead.

MinION read sequences were used to identify the exact CNV copy number for a set of strains. The respective CNV sequences plus 1 kb flanking sequence on either side, were searched against the set of MinION reads for strains MSK478, MSK802, MSK803, MSK812, and UCD321 using BLASTN (v2.10.0) [76]. The search outputs were parsed with RECON-EBB (https://github.com/CMOTsean/recon-ebb) to estimate and visualise copy number from reads which include hits for multiple copies of the CNV sequence and both regions of flanking sequence, i.e the reads which covered the entirety of the repeat region.

### Replication timing profiles

Relative replication time was determined by sort-seq as described previously [77]. Briefly, replicating (S phase) and non-replicating (G2 phase) cells were enriched from an asynchronously growing culture by FACS based on DNA content. In each sample, genomic DNA was extracted and subjected to Illumina sequencing to measure relative DNA copy number. Replication timing profiles were generated by normalizing the replicating (S phase) sample read count to the nonreplicating (G2) sample read count in 1-kb windows.

### Quantitative RT-PCR

Cell harvesting and RNA extraction methods were adapted from Cravener and Mitchell, 2020 [78]. Cells were inoculated from overnight cultures to an A_600_ of 0.2 in 25 ml of pre-warmed YPD broth and incubated at 30°C for 6 h at 200 RPM using an orbital shaker. Cells were then harvested via vacuum filtration using MicroPlus-21 Sterile 0.45μm filters (10407713, Whatman) and stored at −80°C for at least 24 h prior to RNA extraction.Cells were lysed by mechanical disruption as recommended in the QIAGEN RNeasy Mini Kit (74104, QIAGEN) with some modification [78]. Cells were lysed in RLT lysis buffer (QIAGEN) and Phenol:Chloroform:Isoamyl Alcohol (25:24:1)(P3803, Sigma) in a 1:1 ratio. Lysis was performed using a 1600 MiniG from Spex SamplePrep using 30 s lysis followed by 30 s chilling on ice for a total of 6 min. RNA was extracted using a QIAGEN RNeasy Mini Kit followed by two rounds of DNase digestion: one on-column using the QIAGEN RNase-Free DNase Set (79254, QIAGEN); and one off-column using Invitrogen’s TURBO DNA-free Kit (AM1907, ThermoFisher). cDNA was synthesized from 1 μg RNA using M-MLV reverse transcriptase (9PIM170, Promega) and Oligo(dT)15 primers (C110A, Promega), following manufacturer’s instructions. qPCR was performed in 20 μl reactions using 50 ng cDNA using FastStart Universal SYBR Green Master (Rox) (4913850001, Sigma) as per manufacturer’s instructions on an Agilent Technologies Stratagene Mx3005p machine using default “two-step” settings. All primers are listed in S2 Table. Relative quantification was performed using the 2(-Delta Delta C(T)) method by comparing expression to *ACT1*, and using *C. parapsilosis* CLIB214 as the calibrator strain. Calculations and statistics were performed in R using the pcr package [79].

### Data availability

All sequencing data is deposited at NCBI under BioProject number XXXX (awaiting accession numbers).

## Acknowledgements

For Open Access, the authors have applied a CC BY public copyright license to any Author Accepted Manuscript version arising from this submission. This work was supported by Science Foundation Ireland (grant numbers 19/FFP/6668 and 18/CRT/6214 to G.B. and 20/FFP-A/8795 to K.H.W), the Irish Research Council (A.R.), the Chinese Scholarship Scheme (FZ), Deutsche Forschungsgemeinschaft (DFG, German Research Foundation) grant RO-5328/2 (T.R.), National Institutes of Health (NIH) grants R01 AI093808 (T.M.H.), R21 AI156157 (T.M.H.), the Ludwig Center for Cancer Immunotherapy (T.M.H.), the Susan and Peter Solomon Divisional Genomics Program (T.M.H.), and by NIH P30 CA008748 (Cancer Center Core Grant to MSKCC). CAN acknowledges support from the Biotechnology and Biological Sciences Research Council (BBSRC), part of UK Research and Innovation, Core Capability Grant BB/CCG1720/1 and the National Capability (BBS/E/T/000PR9814).

Thanks to the genomics core facility in the Conway Institute UCD for help with Illumina sequencing, to Eoin Ó Cinnéide and Letal Salzberg (UCD) for help with MinION sequencing, and to Lisa Lombardi for help with the adaptive evolution experiment.

---

**Supplementary Table 1** List of strains used.

**Supplementary Table 2.** List of primers used

**Supplementary Figure 1. Copy number determination of CNV-K repeat**

Visualisations of BLASTN results using the CNV-K repeat unit plus 1 kb flanking sequence as query against MinION reads for isolates MSK478 and MSK812 where each plot represents hits against a single read. Each line represents a hit, and adjacent hits are separated vertically for clarity. Read identifiers are shown on the y-axis.

A. The exact copy number of CNV-K at both alleles was identified in isolate MSK812. (i) Seven reads in the MSK812 MinION dataset have 8 copies of the CNV-K repeat unit, one is shown as an example. (ii) Eighteen reads in the MSK812 dataset have 6 copies of the CNV-K repeat unit, one is shown as an example.

B. MSK478 has at least 11 copies on both alleles. No reads in the MSK478 dataset covered the entirety of the repeat array, i.e. no reads had sequence matching both sides of the query flanking DNA. The read with the highest number of copies of the CNV-K repeat contains 11 copies, establishing a likely lower bound for copy number at both alleles.

**Supplementary Figure 2. Structure of CNV-B**

A. (i) CNV-B consists of a central region (B) flanked by two regions (A and C) bounded by inverted repeat pairs (inward-facing triangles). The CNV occurs upstream of the *RTA3* coding sequence. (ii) CNV-B resolves as a repeat array of regions ABC interspersed with inverted copies of region B.

B. The ODIRA model of complex CNV generation, adapted from Brewer et al (2015) [49] under the terms of the Creative Commons Attribution Licence. The topmost diagram has been labelled to demonstrate relationship to the observed CNV in (A).

C. Replication profile of strain MSK802 mapped to the *C. parapsilosis* reference genome. Relative DNA copy number, as a proxy for replication time, is on the Y-axis, where high values denote earlier replication. The region containing *RTA3* on chromosome 1 is denoted by a red bar.

**Supplementary Figure 3. Comparison of Rta2 and Rta3 proteins.**

A. A phylogenetic tree was generated from Muscle alignments of Rta2 and Rta3 sequences from CUG-Ser species (CGOB) using PhyML, implemented in SeaView [55]. Bootstrap values are shown. The pink box highlights the Rta2 and Rta3 sequences from the *Candida parapsilosis* complex. The gene names are taken from the Candida Gene Order Browser.

B. Alignment of *C. parapsilosis* Rta2 and Rta3 generated using Muscle implemented in SeaView.

## References

1. Renne MF, de Kroon AI. The role of phospholipid molecular species in determining the physical properties of yeast membranes. FEBS Lett. 2018;592(8):1330–45.

2. Sharma SC. Implications of sterol structure for membrane lipid composition, fluidity and phospholipid asymmetry in *Saccharomyces cerevisiae*. FEMS Yeast Res. 2006;6(7):1047–51.

3. Grillitsch K, Tarazona P, Klug L, Wriessnegger T, Zellnig G, Leitner E, et al. Isolation and characterization of the plasma membrane from the yeast *Pichia pastoris*. BBA-Biomemb. 2014;1838(7):1889–97.

4. Devaux PF, Herrmann A, Ohlwein N, Kozlov MM. How lipid flippases can modulate membrane structure. BBA-Biomemb. 2008;1778(7-8):1591–600.

5. Andersen JP, Vestergaard AL, Mikkelsen SA, Mogensen LS, Chalat M, Molday RS. P4-ATPases as phospholipid flippases—structure, function, and enigmas. Front Physiol. 2016;7:275.

6. Daleke DL. Regulation of transbilayer plasma membrane phospholipid asymmetry. J Lipid Res. 2003;44(2):233–42.

7. Manente M, Ghislain M. The lipid-translocating exporter family and membrane phospholipid homeostasis in yeast. FEMS Yeast Res. 2009;9(5):673–87.

8. Heimark L, Shipkova P, Greene J, Munayyer H, Yarosh-Tomaine T, DiDomenico B, et al. Mechanism of azole antifungal activity as determined by liquid chromatographic/mass spectrometric monitoring of ergosterol biosynthesis. J Mass Spectrom. 2002;37(3):265–9.

9. Joseph-Horne T, Hollomon DW. Molecular mechanisms of azole resistance in fungi. FEMS Microbiol Lett. 1997;149(2):141–9.

10. Van den Bossche H, Willemsens G, Cools W, Marichal P, Lauwers W. Hypothesis on the molecular basis of the antifungal activity of N-substituted imidazoles and triazoles: Portland Press Ltd.; 1983.

11. Iacano AJ, Lewis H, Hazen JE, Andro H, Smith JD, Gulshan K. Miltefosine increases macrophage cholesterol release and inhibits NLRP3-inflammasome assembly and IL-1β release. Sci Rep. 2019;9(1):1–12.

12. Kaleağasioğlu F, Zaharieva MM, Konstantinov SM, Berger MR. Alkylphospholipids are signal transduction modulators with potential for anticancer therapy. Anticancer Agents Med Chem. 2019;19(1):66–91.

13. Benaim G, Paniz-Mondolfi AE, Sordillo EM, Martinez-Sotillo N. Disruption of intracellular calcium homeostasis as a therapeutic target against *Trypanosoma cruzi*. Front Cell Infect Microbiol. 2020;10:46.

14. Roatt BM, de Oliveira Cardoso JM, De Brito RCF, Coura-Vital W, de Oliveira Aguiar-Soares RD, Reis AB. Recent advances and new strategies on leishmaniasis treatment. Appl Microbiol Biotechnol. 2020:1–13.

15. Papadakis MA, McPhee S, Rabow M. Current medical diagnosis & treatment New York, NY: McGraw-Hill Education LLC; 2018.

16. Dos Reis TF, Horta MAC, Colabardini AC, Fernandes CM, Silva LP, Bastos RW, et al. Screening of chemical llbraries for new antifungal drugs against *Aspergillus fumigatus* reveals sphingolipids are involved in the mechanism of action of miltefosine. mBIO. 2021;12(4):e0145821. Epub 20210810. doi: 10.1128/mBio.01458-21. PubMed PMID: 34372704; PubMed Central PMCID: PMCPMC8406317.

17. Widmer F, Wright LC, Obando D, Handke R, Ganendren R, Ellis DH, et al. Hexadecylphosphocholine (miltefosine) has broad-spectrum fungicidal activity and is efficacious in a mouse model of cryptococcosis. Antimicrob Agents Chemother. 2006;50(2):414–21.

18. Rossi DCP, de Castro Spadari C, Nosanchuk JD, Taborda CP, Ishida K. Miltefosine is fungicidal to *Paracoccidioides* spp. yeast cells but subinhibitory concentrations induce melanisation. Int J Antimicrob Agents. 2017;49(4):465–71.

19. Wu Y, Totten M, Memon W, Ying C, Zhang SX. In vitro antifungal susceptibility of miltefosine alone and in combination with amphotericin B against emerging multidrug resistant *Candida auris*. Antimicrob Agents Chemother. 2019.

20. Barreto TL, Rossato L, de Freitas ALD, Meis JF, Lopes LB, Colombo AL, et al. Miltefosine as an alternative strategy in the treatment of the emerging fungus *Candida auris*. Int J Antimicrob Agents. 2020;56(2):106049.

21. Wu Y, Wu M, Gao J, Ying C. Antifungal activity and mode of action of miltefosine against clinical isolates of *Candida krusei*. Front Microbiol. 2020;11:854.

22. Vila T, Ishida K, Seabra SH, Rozental S. Miltefosine inhibits *Candida albicans* and *non-albicans Candida* spp. biofilms and impairs the dispersion of infectious cells. Int J Antimicrob Agents. 2016;48(5):512–20.

23. Jiménez-López JM, Carrasco MP, Marco C, Segovia JL. Hexadecylphosphocholine disrupts cholesterol homeostasis and induces the accumulation of free cholesterol in HepG2 tumour cells. Biochem Pharmacol. 2006;71(8):1114–21. Epub 20060208. doi: 10.1016/j.bcp.2005.08.001. PubMed PMID: 16466701.

24. Rakotomanga M, Blanc S, Gaudin K, Chaminade P, Loiseau P. Miltefosine affects lipid metabolism in *Leishmania donovani* promastigotes. Antimicrob Agents Chemother. 2007;51(4):1425–30.

25. Pinto-Martinez AK, Rodriguez-Durán J, Serrano-Martin X, Hernandez-Rodriguez V, Benaim G. Mechanism of action of miltefosine on *Leishmania donovani* involves the impairment of acidocalcisome function and the activation of the sphingosine-dependent plasma membrane Ca2+ channel. Antimicrob Agents Chemother. 2018;62(1):e01614–17.

26. Cuesta-Marbán Á, Botet J, Czyz O, Cacharro LM, Gajate C, Hornillos V, et al. Drug uptake, lipid rafts, and vesicle trafficking modulate resistance to an anticancer lysophosphatidylcholine analogue in yeast. J Biol Chem. 2013;288(12):8405–18.

27. Pomorski T, Lombardi R, Riezman H, Devaux PF, van Meer G, Holthuis JC. Drs2p-related P-type ATPases Dnf1p and Dnf2p are required for phospholipid translocation across the yeast plasma membrane and serve a role in endocytosis. Mol Biol Cell. 2003;14(3):1240–54.

28. Hanson PK, Malone L, Birchmore JL, Nichols JW. Lem3p is essential for the uptake and potency of alkylphosphocholine drugs, edelfosine and miltefosine. J Biol Chem. 2003;278(38):36041–50.

29. Pérez-Victoria FJ, Castanys S, Gamarro F. *Leishmania donovani* resistance to miltefosine involves a defective inward translocation of the drug. Antimicrob Agents Chemother. 2003;47(8):2397–403.

30. Pérez-Victoria FJ, Sánchez-Cañete MP, Seifert K, Croft SL, Sundar S, Castanys S, et al. Mechanisms of experimental resistance of *Leishmania* to miltefosine: implications for clinical use. Drug Resistance Updates. 2006;9(1-2):26–39.

31. Seifert K, Matu S, Perez-Victoria FJ, Castanys S, Gamarro F, Croft SL. Characterisation of *Leishmania donovani* promastigotes resistant to hexadecylphosphocholine (miltefosine). Int J Antimicrob Agents. 2003;22(4):380–7.

32. Mondelaers A, Sanchez-Cañete MP, Hendrickx S, Eberhardt E, Garcia-Hernandez R, Lachaud L, et al. Genomic and molecular characterization of miltefosine resistance in *Leishmania infantum* strains with either natural or acquired resistance through experimental selection of intracellular amastigotes. PLoS One. 2016;11(4):e0154101.

33. Srivastava A, Sircaik S, Husain F, Thomas E, Ror S, Rastogi S, et al. Distinct roles of the 7-transmembrane receptor protein Rta3 in regulating the asymmetric distribution of phosphatidylcholine across the plasma membrane and biofilm formation in *Candida albicans*. Cell Microbiol. 2017;19(12). Epub 20171004. doi: 10.1111/cmi.12767. PubMed PMID: 28745020; PubMed Central PMCID: PMCPMC5720375.

34. Kihara A, Igarashi Y. Identification and characterization of a *Saccharomyces cerevisiae* gene, *RSB1*, involved in sphingoid Long-chain Base release. J Biol Chem. 2002;277(33):30048–54.

35. Kihara A, Igarashi Y. Cross talk between sphingolipids and glycerophospholipids in the establishment of plasma membrane asymmetry. Mol Biol Cell. 2004;15(11):4949–59.

36. Johnson SS, Hanson PK, Manoharlal R, Brice SE, Cowart LA, Moye-Rowley WS. Regulation of yeast nutrient permease endocytosis by ATP-binding cassette transporters and a seven-transmembrane protein, RSB1. J Biol Chem. 2010;285(46):35792–802.

37. Soustre I, Letourneux Y, Karst F. Characterization of the *Saccharomyces cerevisiae RTA1* gene involved in 7-aminocholesterol resistance. Curr Genet. 1996;30(2):121–5.

38. Rogers PD, Barker KS. Evaluation of differential gene expression in fluconazole-susceptible and-resistant isolates of *Candida albicans* by cDNA microarray analysis. Antimicrob Agents Chemother. 2002;46(11):3412–7.

39. Rogers PD, Barker KS. Genome-wide expression profile analysis reveals coordinately regulated genes associated with stepwise acquisition of azole resistance in *Candida albicans* clinical isolates. Antimicrob Agents Chemother. 2003;47(4):1220–7.

40. Coste AT, Karababa M, Ischer F, Bille J, Sanglard D. *TAC1,* transcriptional activator of *CDR* genes, is a new transcription factor involved in the regulation of *Candida albicans* ABC transporters *CDR1* and *CDR2*. Eukaryot Cell. 2004;3(6):1639–52.

41. Whaley SG, Tsao S, Weber S, Zhang Q, Barker KS, Raymond M, et al. The *RTA3* gene, encoding a putative lipid translocase, influences the susceptibility of *Candida albicans* to fluconazole. Antimicrob Agents Chemother. 2016;60(10):6060–6.

42. Zhai B, Ola M, Rolling T, Tosini NL, Joshowitz S, Littmann ER, et al. High-resolution mycobiota analysis reveals dynamic intestinal translocation preceding invasive candidiasis. Nat Med. 2020;26(1):59–64.

43. West PT, Peters SL, Olm MR, Feiqiao BY, Gause H, Lou YC, et al. Genetic and behavioral adaptation of *Candida parapsilosis* to the microbiome of hospitalized infants revealed by *in situ* genomics, transcriptomics, and proteomics. Microbiome. 2021;9(1):1–17.

44. McKenna A, Hanna M, Banks E, Sivachenko A, Cibulskis K, Kernytsky A, et al. The Genome Analysis Toolkit: a MapReduce framework for analyzing next-generation DNA sequencing data. Genome Res. 2010;20(9):1297–303.

45. Lischer HE, Excoffier L, Heckel G. Ignoring heterozygous sites biases phylogenomic estimates of divergence times: implications for the evolutionary history of Microtus voles. Mol Biol Evol. 2014;31(4):817–31.

46. Asadzadeh M, Dashti M, Ahmad S, Alfouzan W, Alameer A. Whole-Genome and Targeted-Amplicon sequencing of fluconazole-susceptible and-resistant *Candida parapsilosis* isolates from Kuwait reveals a previously undescribed N1132D polymorphism in CDR1. Antimicrob Agents Chemother. 2021;65(2):e01633–20.

47. Lopez-Delisle L, Rabbani L, Wolff J, Bhardwaj V, Backofen R, Grüning B, et al. pyGenomeTracks: reproducible plots for multivariate genomic datasets. Bioinformatics. 2021;37(3):422.

48. Carvalho CM, Zhang F, Liu P, Patel A, Sahoo T, Bacino CA, et al. Complex rearrangements in patients with duplications of MECP2 can occur by fork stalling and template switching. Hum Mol Genet. 2009;18(12):2188–203.

49. Brewer BJ, Payen C, Di Rienzi SC, Higgins MM, Ong G, Dunham MJ, et al. Origin-dependent inverted-repeat amplification: tests of a model for inverted DNA amplification. PLoS Genet. 2015;11(12):e1005699.

50. Müller CA, Hawkins M, Retkute R, Malla S, Wilson R, Blythe MJ, et al. The dynamics of genome replication using deep sequencing. Nucleic Acids Res. 2014;42(1):e3–e.

51. Ropars J, Maufrais C, Diogo D, Marcet-Houben M, Perin A, Sertour N, et al. Gene flow contributes to diversification of the major fungal pathogen *Candida albicans*. Nat Commun. 2018;9(1):2253. Epub 20180608. doi: 10.1038/s41467-018-04787-4. PubMed PMID: 29884848; PubMed Central PMCID: PMCPMC5993739.

52. Sitterlé E, Maufrais C, Sertour N, Palayret M, d’Enfert C, Bougnoux ME. Within-host genomic diversity of *Candida albicans* in healthy carriers. Sci Rep. 2019;9(1):2563. Epub 20190222. doi: 10.1038/s41598-019-38768-4. PubMed PMID: 30796326; PubMed Central PMCID: PMCPMC6385308.

53. Lombardi L, Oliveira-Pacheco J, Butler G. Plasmid-based CRISPR-Cas9 gene editing in multiple *Candida* species. mSphere. 2019;4(2):e00125–19.

54. Krogh A, Larsson B, Von Heijne G, Sonnhammer EL. Predicting transmembrane protein topology with a hidden Markov model: application to complete genomes. J Mol Biol. 2001;305(3):567–80.

55. Gouy M, Guindon S, Gascuel O. SeaView version 4: a multiplatform graphical user interface for sequence alignment and phylogenetic tree building. Mol Biol Evol. 2010;27(2):221–4.

56. Van der Mark VA, Elferink RP, Paulusma CC. P4 ATPases: flippases in health and disease. Int J Mol Sci. 2013;14(4):7897–922.

57. Makuta H, Obara K, Kihara A. Loop 5 region is important for the activity of the long-chain base transporter Rsb1. The journal of biochemistry. 2017;161(2):207–13.

58. Fitzpatrick DA, O’Gaora P, Byrne KP, Butler G. Analysis of gene evolution and metabolic pathways using the *Candida* Gene Order Browser. BMC Genomics. 2010;11(1):1–14.

59. Maguire SL, ÓhÉigeartaigh SS, Byrne KP, Schröder MS, O’Gaora P, Wolfe KH, et al. Comparative genome analysis and gene finding in *Candida* species using CGOB. Mol Biol Evol. 2013;30(6):1281–91.

60. Jia X, Wang Y, Jia Y, Gao P, Xu Y, Wang L, et al. *RTA2* is involved in calcineurin-mediated azole resistance and sphingoid long-chain base release in *Candida albicans*. Cell Mol Life Sci. 2009;66(1):122–34.

61. Thomas E, Sircaik S, Roman E, Brunel J-M, Johri AK, Pla J, et al. The activity of *RTA2*, a downstream effector of the calcineurin pathway, is required during tunicamycin-induced ER stress response in *Candida albicans*. FEMS Yeast Res. 2015;15(8).

62. Yang F, Gritsenko V, Slor Futterman Y, Gao L, Zhen C, Lu H, et al. Tunicamycin potentiates antifungal drug tolerance via aneuploidy in *Candida albicans*. mBIO. 2021;12(4):e02272–21.

63. Sakofsky CJ, Ayyar S, Deem AK, Chung W-H, Ira G, Malkova A. Translesion polymerases drive microhomology-mediated break-induced replication leading to complex chromosomal rearrangements. Mol Cell. 2015;60(6):860–72.

64. Fogel S, Welch JW. Tandem gene amplification mediates copper resistance in yeast. Proc Natl Acad Sci (USA). 1982;79(17):5342–6.

65. Hull RM, Cruz C, Jack CV, Houseley J. Environmental change drives accelerated adaptation through stimulated copy number variation. PLoS Biol. 2017;15(6):e2001333.

66. Todd RT, Selmecki A. Expandable and reversible copy number amplification drives rapid adaptation to antifungal drugs. Elife. 2020;9:e58349.

67. Papp C, Kocsis K, Tóth R, Bodai L, Willis JR, Ksiezopolska E, et al. Echinocandin-induced microevolution of *Candida parapsilosis* influences virulence and abiotic stress tolerance. mSphere. 2018;3(6):e00547–18.

68. Ene IV, Farrer RA, Hirakawa MP, Agwamba K, Cuomo CA, Bennett RJ. Global analysis of mutations driving microevolution of a heterozygous diploid fungal pathogen. Proc Natl Acad Sci (USA). 2018;115(37):E8688–E97.

69. De Coster W, D’Hert S, Schultz DT, Cruts M, Van Broeckhoven C. NanoPack: visualizing and processing long-read sequencing data. Bioinformatics. 2018;34(15):2666–9.

70. Jiang H, Lei R, Ding S-W, Zhu S. Skewer: a fast and accurate adapter trimmer for next-generation sequencing paired-end reads. BMC Bioinformatics. 2014;15(1):1–12.

71. Vaser R, Adusumalli S, Leng SN, Sikic M, Ng PC. SIFT missense predictions for genomes. Nat Protoc. 2016;11(1):1–9.

72. Suzek BE, Wang Y, Huang H, McGarvey PB, Wu CH, Consortium U. UniRef clusters: a comprehensive and scalable alternative for improving sequence similarity searches. Bioinformatics. 2015;31(6):926–32.

73. Thorvaldsdóttir H, Robinson JT, Mesirov JP. Integrative Genomics Viewer (IGV): high-performance genomics data visualization and exploration. Brief Bioinform. 2013;14(2):178–92.

74. Stamatakis A. RAxML version 8: a tool for phylogenetic analysis and post-analysis of large phylogenies. Bioinformatics. 2014;30(9):1312–3.

75. Quinlan AR, Hall IM. BEDTools: a flexible suite of utilities for comparing genomic features. Bioinformatics. 2010;26(6):841–2.

76. Altschul SF, Madden TL, Schäffer AA, Zhang J, Zhang Z, Miller W, et al. Gapped BLAST and PSI-BLAST: a new generation of protein database search programs. Nucleic Acids Res. 1997;25(17):3389–402.

77. Batrakou DG, Müller CA, Wilson RHC, Nieduszynski CA. DNA copy-number measurement of genome replication dynamics by high-throughput sequencing: the sort-seq, sync-seq and MFA-seq family. Nat Protoc. 2020;15(3):1255–84. Epub 20200212. doi: 10.1038/s41596-019-0287-7. PubMed PMID: 32051615.

78. Cravener MV, Mitchell AP. *Candida albicans* culture, cell Harvesting, and total RNA extraction. Bio-protocol. 2020;10(21):e3803–e.

79. Ahmed M, Kim DR. pcr: an R package for quality assessment, analysis and testing of qPCR data. PeerJ. 2018;6:e4473.

