## Supplementary material for "Resistance to miltefosine results from amplification of the *RTA3* floppase or inactivation of flippases in *Candida parapsilosis*"

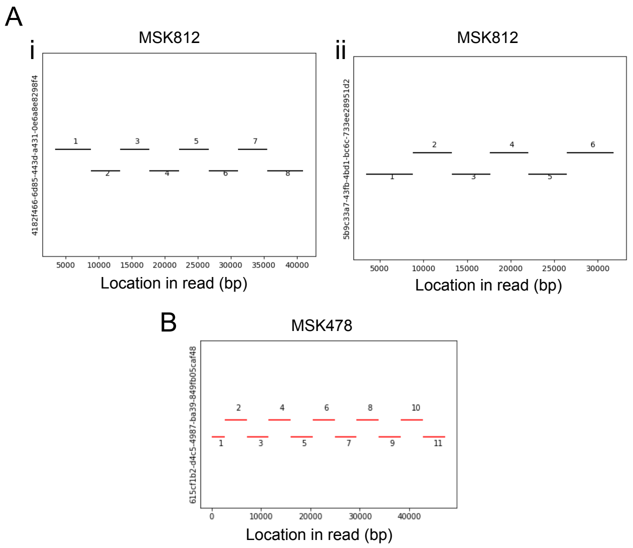

**Supplementary Figure 1 Copy number determination of CNV-K repeat.**

Visualisations of BLASTN results using the CNV-K repeat unit plus 1 kb flanking sequence as query against MinION reads for isolates MSK478 and MSK812 where each plot represents hits against a single read. Each line represents a hit, and adjacent hits are separated vertically for clarity. Read identifiers are shown on the y-axis.

A. The exact copy number of CNV-K at both alleles was identified in isolate MSK812. (i) Seven reads in the MSK812 MinION dataset have 8 copies of the CNV-K repeat unit, one is shown as an example. (ii) Eighteen reads in the MSK812 dataset have 6 copies of the CNV-K repeat unit, one is shown as an example.

B. MSK478 has at least 11 copies on both alleles. No reads in the MSK478 data-set covered the entirety of the repeat array, i.e. no reads had sequence matching both sides of the query flanking DNA. The read with the highest number of copies of the CNV-K repeat contains 11 copies, establishing a likely lower bound for copy number at both alleles.

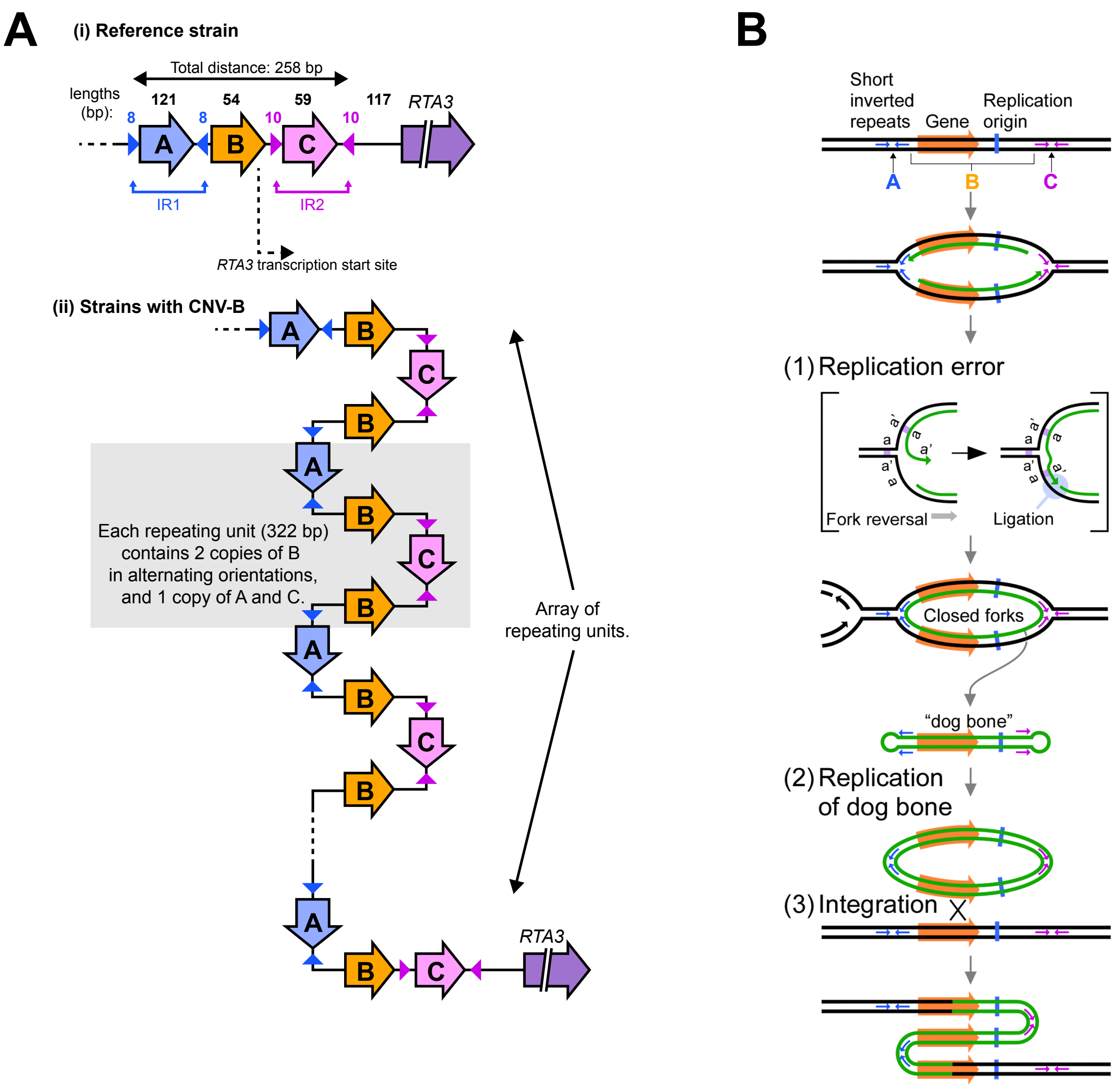

### Supplementary Figure 2. Structure of CNV-B

A. (i) CNV-B consists of a central region (B) flanked by two regions (A and C) bounded by inverted repeat pairs (inward-facing triangles). The CNV occurs upstream of the RTA3 coding sequence. (ii) CNV-B resolves as a repeat array of regions ABC interspersed with inverted copies of region B.

B. The ODIRA model of complex CNV generation, adapted from Brewer et al (2015) [49] under the terms of the Creative Commons Attribution Licence. The topmost diagram has been labelled to demonstrate relationship to the observed CNV in (A).

C. (overleaf) Replication profile of strain MSK802 mapped to the *C. parapsilosis* reference genome. Relative DNA copy number, as a proxy for replication time, is on the Y-axis, where high values denote earlier replication. The region containing *RTA3* on chromosome 1 is denoted by a red bar.

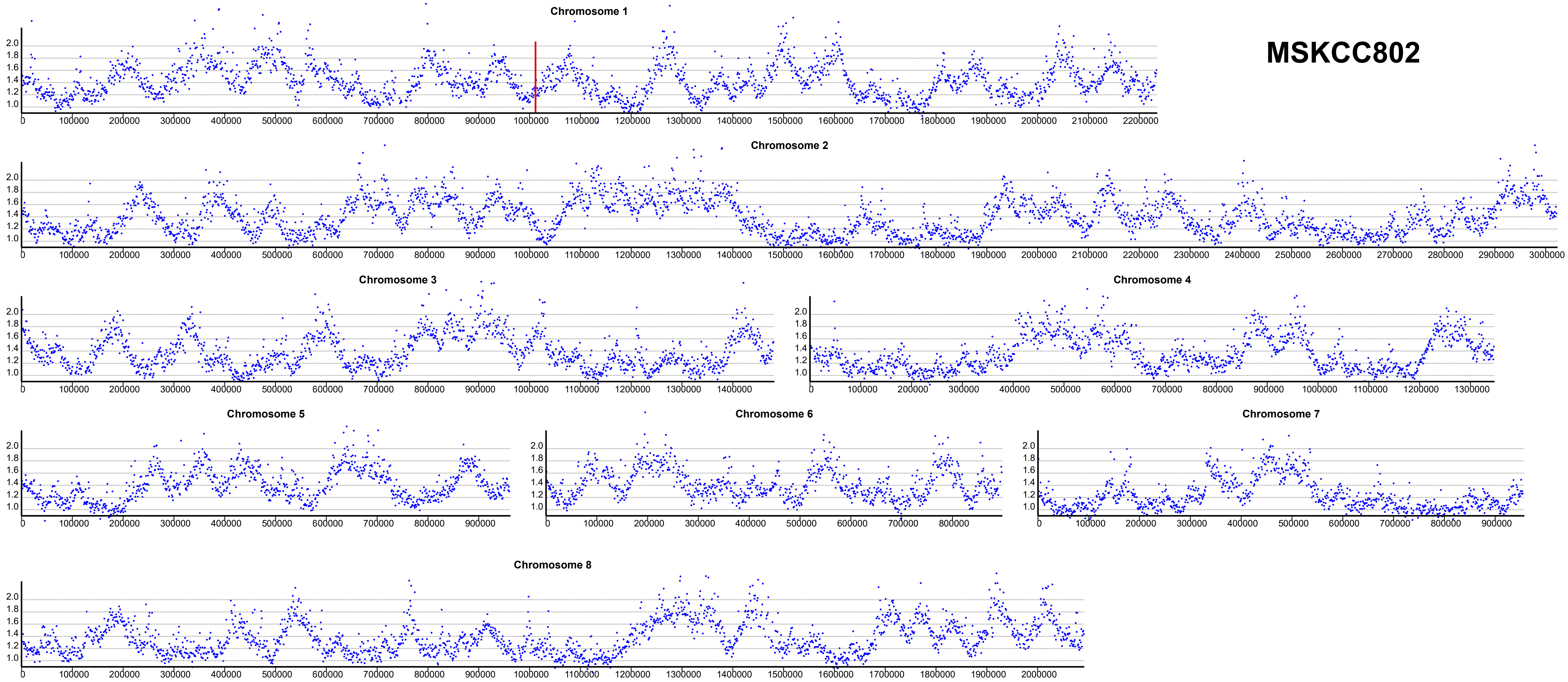
MSKCC802
Chromosome 1
Chromosome 2
Chromosome 3
Chromosome 4
Chromosome 5
Chromosome 6
Chromosome 7
Chromosome 8

Table S1. List of strains

| Strain | Location[1] | Species | CNV | Copy No. | Clade | Data Source |
|---|---|---|---|---|---|---|
| 103 | UK | *C. parapsilosis* | - | 2 | 5 | This study |
| 02-203 | Italy | *C. parapsilosis* | - | 2 | 5 | This study |
| 73-037 | Leeds, UK | *C. parapsilosis* | - | 2 | 2 | This study |
| 73-107 | London, UK | *C. parapsilosis* | - | 2 | 2 | This study |
| 73-114 | Leeds, UK | *C. parapsilosis* | - | 2 | 5 | This study |
| 74-046 | Leeds, UK | *C. parapsilosis* | - | 2 | 5 | This study |
| 81-040 | UK | *C. parapsilosis* | - | 2 | 5 | This study |
| 81-042 | Leeds, UK | *C. parapsilosis* | - | 2 | 3 | This study |
| 81-253 | UK | *C. parapsilosis* | - | 2 | 5 | This study |
| 90-137 | USA | *C. parapsilosis* | - | 2 | 5 | PRJNA563885 |
| BC014S | Portugal | *C. parapsilosis* | - | 2 | 1 | PRJNA326748 |
| CBS1954 | Italy | *C. parapsilosis* | - | 2 | 2 | PRJEB1831 |
| CBS6318 | USA | *C. parapsilosis* | - | 2 | 3 | PRJEB1831 |
| CDC173 | Mississippi, USA | *C. parapsilosis* | - | 2 | 4 | This study |
| CDC179 | Mississippi, USA | *C. parapsilosis* | - | 2 | 4 | This study |
| CDC317 | Mississippi, USA | *C. parapsilosis* | - | 2 | 4 | This study |
| CLIB214 | Puerto Rico | *C. parapsilosis* | - | 2 | 2 | PRJNA563885 |
| CP35176 | Unknown | *C. parapsilosis* | - | 2 | 2 | PRJNA361149 |
| CP35177 | Unknown | *C. parapsilosis* | - | 2 | 2 | PRJNA361149 |
| GA1 | Germany | *C. parapsilosis* | - | 2 | 4 | PRJEB1831 |
| J931058 | Belgium | *C. parapsilosis* | - | 2 | 5 | This study |
| J931845 | Japan | *C. parapsilosis* | - | 2 | 5 | This study |
| J950218 | USA | *C. parapsilosis* | - | 2 | 3 | This study |
| J951066 | Korea | *C. parapsilosis* | - | 2 | 4 | This study |
| J961250 | Lisbon, Portugal | *C. parapsilosis* | - | 2 | 5 | This study |
| UCD321 | Ireland | *C. parapsilosis* | D | 27 | 5 | This study |
| yHMJ4 | USA | *C. parapsilosis* | - | 2 | 1 | This study |
| YV1 | Tuscany | *C. parapsilosis* | J | 45 | 4 | PRJEB1831 |
| Kw1590-18 | Kuwait | *C. parapsilosis* | - | 2 | 4 | PRJNA661583 |
| Kw2006-15 | Kuwait | *C. parapsilosis* | - | 2 | 5 | PRJNA661583 |
| Kw3259-15 | Kuwait | *C. parapsilosis* | Z | 24 | 1 | PRJNA661583 |

A

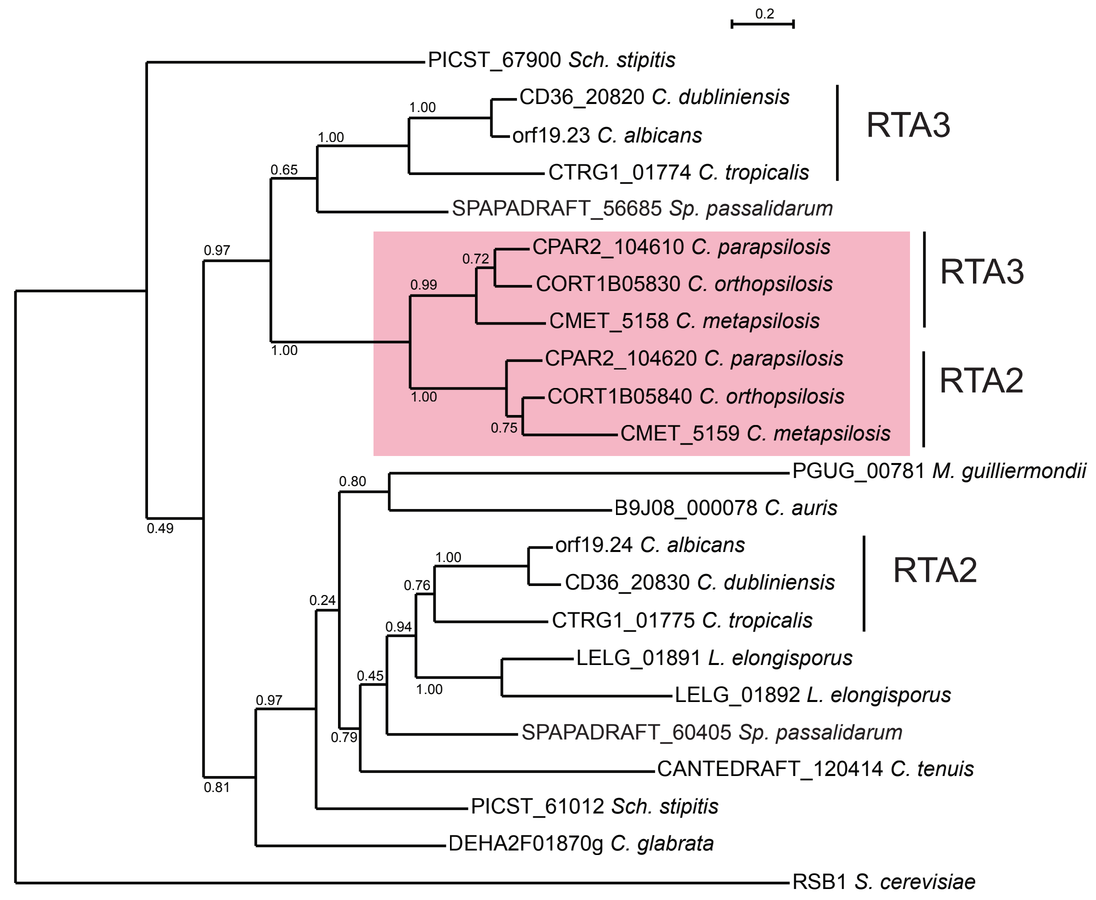

B

```
       1
RTA2   MDQSPSIAAW TP--TTTATS TSLKSVATSA HSALESSLSK ALHSLTGHTT TMERNDYIHA SRAARGAQAS
RTA3   MDQLASITAW TPSSTNTPTT TSLKSVATTA QSALESSLSA ALRDFSSYGT KINNEQYVSA SRAIRGAQAS

      71
RTA2   LSIISGENIL ATATAESIKA QATEMIWNST QNLEDLAWEE NMYGIRRLTR PGNIFFLAAF SITFLWYTLI
RTA3   LSIISAEHVL ATATADNIKA QATELIWNST QNLDDLWWDE NVYDIRRLTR PGNIIFLVVF AISFLWYIVI

     141
RTA2   FIKSRYWWFN IAFFCGALLE FLGFLGRVLS FNDMSSFSYY VMQMVTLTIA PAFTMGGIYF LLGQLVIIHG
RTA3   LAKSRYWWFN VAFFCGAALE FLGFLGRVLS FNDMTYFPYY LLQLIVLTIA PAFIMGGIYF LLGQLVIIHG

     211
RTA2   RQFSFLKPLW YAYIFIACDI LSLFIQAAGG GVASAATSGN TSPDPGTNTM IAGIAFQAFA MSIYVSLWFV
RTA3   RQFSVLKPLW YAYIFIACDI TSLVIQAIGG GIASVRAENY ENADPGTYTM IAGIAFQVFS MSIYIIFWTI

     281
RTA2   FIWKIYFPKK DTH-ARYAIE KD---NQQQM LKPGLKNFLK LFFNTKASAE YKRTVLEPFY NPKYSDIRQR
RTA3   FLWKIYFPKT DKYPSQLASE KSDLVDPQQV LKPSVKNFFK LLINSKKTAE YKRNILEPYY NPKYADIRSR

     351
RTA2   PLYNYFPLAV TCAIIAIFIR CIYRVVELAQ GFRGYLITHE VYIMCLDAAM LVIALYVFIP FHPYIVFGSS
RTA3   KMYDYFPLAV SLAIIAVFVR CIYRVVELAQ GFSGYLITHE VYIMCLDALM LFIALYMFIP FHPHIVFGAS

     421
RTA2   NIIRLGDIKG NADEKLEED- ---------- ---------- ---------- ---------- ----------
RTA3   NIIRLGDIKG NTDEKIEEEV LYPVDDGYDS NATRKGTAVG SPYGETQDLE KHGSETSYKA DLGDANHQAS

     491
RTA2   ---ETGLTKM DNNTSKSESI --EPVLMDSI EK*------- ---
RTA3   YPIESDYTPQ GIHSANNSHI PRDSYPLESR EEYHDAYGTR HR*
```

**Supplementary Figure 3. Comparison of Rta2 and Rta3 proteins.**

(A) A phylogenetic tree was generated from Muscle alignments of Rta2 and Rta3 sequences from CUG-Ser species (CGOB) using PhyML, implemented in SeaView [55]. Bootstrap values are shown. The pink box highlights the Rta2 and Rta3 sequences from the Candida parapsilosis complex. The gene names are taken from the Candida Gene Order Browser.

(B) Alignment of C. parapsilosis Rta2 and Rta3 generated using Muscle implemented in SeaView.

| | | | | | | |
|---|---|---|---|---|---|---|
| FM02 | Nantes, France | *C. parapsilosis* | N | 8 | 4 | This study |
| FM03 | Nantes, France | *C. parapsilosis* | N | 8 | 4 | This study |
| FM05 | Nantes, France | *C. parapsilosis* | N | 8 | 4 | This study |
| FM06 | Nantes, France | *C. parapsilosis* | F | 5 | 1 | This study |
| FM07 | Nantes, France | *C. parapsilosis* | D | 16 | 5 | This study |
| FM10 | Nantes, France | *C. parapsilosis* | N | 13 | 4 | This study |
| FM13 | Nantes, France | *C. parapsilosis* | O | 13 | 4 | This study |
| FM14 | Nantes, France | *C. parapsilosis* | - | 2 | 2 | This study |
| FM16 | Nantes, France | *C. parapsilosis* | J | 50 | 4 | This study |
| FM17 | Nantes, France | *C. parapsilosis* | N | 8 | 4 | This study |
| FM20 | Nantes, France | *C. parapsilosis* | N | 8 | 4 | This study |
| FM21 | Nantes, France | *C. parapsilosis* | D | 24 | 5 | This study |
| FM32 | Nantes, France | *C. parapsilosis* | L | 19 | 3 | This study |
| FM36 | Nantes, France | *C. parapsilosis* | D | 24 | 5 | This study |
| FM41 | Nantes, France | *C. parapsilosis* | L | 12 | 3 | This study |
| FM43 | Nantes, France | *C. parapsilosis* | F | 15 | 1 | This study |
| MSK1 | New York | *C. parapsilosis* | K | 29 | 1 | PRJNA579121 |
| MSK1004 | New York | *C. parapsilosis* | K | 24 | 1 | This study |
| MSK1015 | New York | *C. parapsilosis* | K | 24 | 1 | This study |
| MSK1082 | New York (Pt11) | *C. parapsilosis* | D | 11 | 5 | This study |
| MSK1090 | New York (Pt11) | *C. parapsilosis* | D | 11 | 5 | This study |
| MSK1119 | New York (Pt1) | *C. parapsilosis* | K | 13 | 1 | This study |
| MSK1129 | New York (Pt1) | *C. parapsilosis* | K | 13 | 1 | This study |
| MSK1191 | New York (Pt2) | *C. parapsilosis* | - | 2 | 1 | This study |
| MSK1206 | New York | *C. parapsilosis* | F | 7 | 1 | This study |
| MSK1241 | New York (Pt10) | *C. parapsilosis* | D | 17 | 5 | This study |
| MSK1258 | New York (Pt10) | *C. parapsilosis* | D | 17 | 5 | This study |
| MSK1286 | New York | *C. parapsilosis* | - | 2 | 4 | This study |
| MSK1298 | New York | *C. parapsilosis* | - | 2 | 4 | This study |
| MSK1302 | New York | *C. parapsilosis* | - | 2 | 4 | This study |
| MSK1324 (Pt9) | New York | *C. parapsilosis* | A | 6 | 5 | This study |
| MSK1325 | New York (Pt9) | *C. parapsilosis* | A | 6 | 5 | This study |
| MSK1351 | New York | *C. parapsilosis* | - | 2 | 4 | This study |

| | | | | | | |
|---|---|---|---|---|---|---|
| MSK1394 | New York (Pt5) | *C. parapsilosis* | F | 7 | 1 | This study |
| MSK1398 | New York (Pt5) | *C. parapsilosis* | F | 7 | 1 | This study |
| MSK1399 | New York (Pt5) | *C. parapsilosis* | F | 7 | 1 | This study |
| MSK158 | New York | *C. parapsilosis* | - | 2 | 4 | This study |
| MSK1617 | New York | *C. parapsilosis* | - | 2 | 1 | This study |
| MSK1666 | New York | *C. parapsilosis* | - | 2 | 1 | This study |
| MSK17 | New York | *C. parapsilosis* | K | 29 | 1 | PRJNA579121 |
| MSK1700 | New York (Pt9) | *C. parapsilosis* | A | 6 | 5 | This study |
| MSK1762 | New York | *C. parapsilosis* | K | 26 | 1 | This study |
| MSK18 | New York | *C. parapsilosis* | K | 27 | 1 | PRJNA579121 |
| MSK19 | New York | *C. parapsilosis* | K | 29 | 1 | PRJNA579121 |
| MSK2 | New York | *C. parapsilosis* | K | 29 | 1 | PRJNA579121 |
| MSK2049 | New York | *C. parapsilosis* | K | 27 | 1 | This study |
| MSK2057 | New York | *C. parapsilosis* | K | 27 | 1 | This study |
| MSK2060 | New York | *C. parapsilosis* | K | 28 | 1 | This study |
| MSK2084 | New York | *C. parapsilosis* | K | 27 | 1 | This study |
| MSK2086 | New York | *C. parapsilosis* | K | 27 | 1 | This study |
| MSK2092 | New York | *C. parapsilosis* | K | 27 | 1 | This study |
| MSK2094 | New York | *C. parapsilosis* | K | 27 | 1 | This study |
| MSK2107 | New York (Pt6) | *C. parapsilosis* | A | 9 | 1 | This study |
| MSK2108 | New York (Pt6) | *C. parapsilosis* | A | 9 | 1 | This study |
| MSK2123 | New York (Pt7) | *C. parapsilosis* | - | 2 | 2 | This study |
| MSK2124 | New York (Pt7) | *C. parapsilosis* | - | 2 | 2 | This study |
| MSK2131 | New York | *C. parapsilosis* | K | 28 | 1 | This study |
| MSK2134 | New York | *C. parapsilosis* | K | 29 | 1 | This study |
| MSK2141 | New York | *C. parapsilosis* | K | 27 | 1 | This study |
| MSK2159 | New York (Pt3) | *C. parapsilosis* | F | 5 | 1 | This study |
| MSK2160 | New York | *C. parapsilosis* | K | 27 | 1 | This study |
| MSK2161 | New York (Pt3) | *C. parapsilosis* | F | 5 | 1 | This study |
| MSK2162 | New York | *C. parapsilosis* | K | 27 | 1 | This study |
| MSK2191 | New York | *C. parapsilosis* | - | 2 | 4 | This study |
| MSK2199 | New York | *C. parapsilosis* | A | 15 | 5 | This study |
| MSK2233 | New York (Pt8) | *C. parapsilosis* | - | 2 | 2 | This study |

| | | | | | | |
|---|---|---|---|---|---|---|
| MSK2234 | New York (Pt8) | *C. parapsilosis* | - | 2 | 2 | This study |
| MSK2248 | New York | *C. parapsilosis* | K | 27 | 1 | This study |
| MSK2384 | New York | *C. parapsilosis* | L | 14 | 3 | This study |
| MSK2386 | New York | *C. parapsilosis* | - | 2 | 2 | This study |
| MSK2387 | New York | *C. parapsilosis* | B | 22[2] | 1 | This study |
| MSK2389 | New York | *C. parapsilosis* | B | 13[2] | 1 | This study |
| MSK2390 | New York | *C. parapsilosis* | - | 2 | 1 | This study |
| MSK247 | New York | *C. parapsilosis* | - | 2 | 5 | PRJNA579121 |
| MSK249 | New York | *C. parapsilosis* | - | 2 | 4 | PRJNA579121 |
| MSK250 | New York | *C. parapsilosis* | - | 2 | 4 | PRJNA579121 |
| MSK251 | New York | *C. parapsilosis* | - | 2 | 4 | PRJNA579121 |
| MSK264 | New York | *C. parapsilosis* | K | 25 | 1 | PRJNA579121 |
| MSK265 | New York | *C. parapsilosis* | - | 2 | 4 | PRJNA579121 |
| MSK266 | New York | *C. parapsilosis* | - | 2 | 4 | PRJNA579121 |
| MSK281 | New York | *C. parapsilosis* | - | 2 | 4 | PRJNA579121 |
| MSK282 | New York | *C. parapsilosis* | K | 28 | 1 | PRJNA579121 |
| MSK283 | New York | *C. parapsilosis* | - | 2 | 4 | PRJNA579121 |
| MSK296 | New York | *C. parapsilosis* | K | 31 | 1 | PRJNA579121 |
| MSK297 | New York | *C. parapsilosis* | K | 29 | 1 | PRJNA579121 |
| MSK298 | New York | *C. parapsilosis* | K | 29 | 1 | PRJNA579121 |
| MSK313 | New York | *C. parapsilosis* | K | 27 | 1 | PRJNA579121 |
| MSK314 | New York | *C. parapsilosis* | K | 34 | 1 | PRJNA579121 |
| MSK315 | New York | *C. parapsilosis* | K | 34 | 1 | PRJNA579121 |
| MSK33 | New York | *C. parapsilosis* | K | 26 | 1 | PRJNA579121 |
| MSK34 | New York | *C. parapsilosis* | K | 26 | 1 | PRJNA579121 |
| MSK35 | New York | *C. parapsilosis* | K | 29 | 1 | PRJNA579121 |
| MSK478 | New York | *C. parapsilosis* | K | 25 | 1 | PRJNA579121 |
| MSK485 | New York | *C. parapsilosis* | K | 25 | 1 | This study |
| MSK486 | New York | *C. parapsilosis* | K | 29 | 1 | This study |
| MSK489 | New York (Pt4) | *C. parapsilosis* | F | 7 | 1 | This study |
| MSK49 | New York | *C. parapsilosis* | K | 29 | 1 | PRJNA579121 |
| MSK490 | New York (Pt4) | *C. parapsilosis* | F | 8 | 1 | This study |
| MSK5 | New York | *C. parapsilosis* | K | 29 | 1 | PRJNA579121 |

| | | | | | | |
|---|---|---|---|---|---|---|
| MSK51 | New York | *C. parapsilosis* | K | 25 | 1 | PRJNA579121 |
| MSK519 | New York | *C. parapsilosis* | K | 25 | 1 | This study |
| MSK52 | New York | *C. parapsilosis* | K | 27 | 1 | PRJNA579121 |
| MSK520 | New York | *C. parapsilosis* | K | 26 | 1 | This study |
| MSK522 | New York | *C. parapsilosis* | K | 25 | 1 | This study |
| MSK536 | New York | *C. parapsilosis* | K | 27 | 1 | This study |
| MSK543 | New York | *C. parapsilosis* | K | 26 | 1 | This study |
| MSK544 | New York | *C. parapsilosis* | K | 23 | 1 | This study |
| MSK608 | New York | *C. parapsilosis* | - | 2 | 4 | This study |
| MSK611 | New York | *C. parapsilosis* | - | 2 | 4 | This study |
| MSK612 | New York | *C. parapsilosis* | - | 2 | 4 | This study |
| MSK620 | New York (Pt2) | *C. parapsilosis* | - | 2 | 1 | This study |
| MSK624 | New York | *C. parapsilosis* | K | 26 | 1 | This study |
| MSK630 | New York | *C. parapsilosis* | K | 26 | 1 | This study |
| MSK65 | New York | *C. parapsilosis* | K | 27 | 1 | PRJNA579121 |
| MSK67 | New York | *C. parapsilosis* | K | 29 | 1 | PRJNA579121 |
| MSK794 | New York (Pt3) | *C. parapsilosis* | F | 5 | 1 | This study |
| MSK795 | New York | *C. parapsilosis* | - | 2 | 1 | This study |
| MSK799 | New York | *C. parapsilosis* | E | 19 | 1 | This study |
| MSK800 | New York | *C. parapsilosis* | C | 29 | 1 | This study |
| MSK802 | New York | *C. parapsilosis* | B | 27[2] | 1 | This study |
| MSK803 | New York | *C. parapsilosis* | B | 24[2] | 1 | This study |
| MSK804 | New York | *C. parapsilosis* | - | 2 | 4 | This study |
| MSK806 | New York | *C. parapsilosis* | F | 10 | 1 | This study |
| MSK807 | New York | *C. parapsilosis* | G | 4 | 5 | This study |
| MSK808 | New York | *C. parapsilosis* | I | 42 | 1 | This study |
| MSK809 | New York | *C. parapsilosis* | A | 14 | 5 | This study |
| MSK810 | New York | *C. parapsilosis* | H | 19 | 1 | This study |
| MSK811 | New York | *C. parapsilosis* | - | 2 | 2 | This study |
| MSK812 | New York | *C. parapsilosis* | K | 14 | 1 | This study |
| MSK813 | New York | *C. parapsilosis* | C | 31 | 1 | This study |
| MSK814 | New York | *C. parapsilosis* | - | 2 | 3 | This study |
| MSK815 | New York | *C. parapsilosis* | D | 17 | 5 | This study |

| | | | | | | |
|---|---|---|---|---|---|---|
| MSK844 | New York | *C. parapsilosis* | - | 2 | 2 | This study |
| MSK846 | New York | *C. parapsilosis* | M | 6 | 5 | This study |
| MSK848 | New York | *C. parapsilosis* | - | 2 | 4 | This study |
| MSK850 | New York | *C. parapsilosis* | - | 2 | 4 | This study |
| MSK891 | New York | *C. parapsilosis* | - | 2 | 4 | This study |
| MSK923 | New York | *C. parapsilosis* | - | 2 | 4 | This study |
| MSK925 | New York | *C. parapsilosis* | - | 2 | 4 | This study |
| 151 | Atlanta, USA | *C. orthopsilosis* | Co2 | 13 | N/A | PRJNA322245 |
| 1540 | Baltimore, USA | *C. orthopsilosis* | Co3 | 19 | N/A | PRJNA322245 |
| 1799 | Atlanta, USA | *C. orthopsilosis* | Co2 | 24 | N/A | PRJNA322245 |
| 1825 | Baltimore, USA | *C. orthopsilosis* | Co2 | 15 | N/A | PRJNA322245 |
| 185 | Atlanta, USA | *C. orthopsilosis* | Co2 | 13 | N/A | PRJNA322245 |
| 282 | Baltimore, USA | *C. orthopsilosis* | - | 2 | N/A | PRJNA322245 |
| 320 | Baltimore, USA | *C. orthopsilosis* | - | 2 | N/A | PRJNA322245 |
| 421 | Pisa, Italy | *C. orthopsilosis* | Co2 | 19 | N/A | PRJNA322245 |
| 422 | Pisa, Italy | *C. orthopsilosis* | Co3 | 21 | N/A | PRJNA322245 |
| 423 | L'Aquila, Italy | *C. orthopsilosis* | Co2 | 39 | N/A | PRJNA322245 |
| 424 | Pisa, Italy | *C. orthopsilosis* | - | 2 | N/A | PRJNA322245 |
| 425 | Pisa, Italy | *C. orthopsilosis* | Co3 | 18 | N/A | PRJNA322245 |
| 426 | Varese, Italy | *C. orthopsilosis* | - | 2 | N/A | PRJNA322245 |
| 427 | Pisa, Italy | *C. orthopsilosis* | - | 2 | N/A | PRJNA322245 |
| 428 | Hong Kong | *C. orthopsilosis* | - | 2 | N/A | PRJNA322245 |
| 433 | St. Niklaas, Belgium | *C. orthopsilosis* | - | 2 | N/A | PRJNA322245 |
| 434 | NCPF, UK | *C. orthopsilosis* | Co4 | 1 | N/A | PRJNA322245 |
| 435 | Pisa, Italy | *C. orthopsilosis* | - | 2 | N/A | PRJNA322245 |
| 436 | Pisa, Italy | *C. orthopsilosis* | - | 2 | N/A | PRJNA322245 |
| 437 | Pisa, Italy | *C. orthopsilosis* | Co3 | 19 | N/A | PRJNA322245 |
| MSK477 | New York | *C. orthopsilosis* | Co1 | 8 | N/A | PRJNA579121 |
| MSK479 | New York | *C. orthopsilosis* | Co1 | 9 | N/A | PRJNA579121 |
| 498 | Baltimore, USA | *C. orthopsilosis* | - | 2 | N/A | PRJNA322245 |
| 504 | Pisa, Italy | *C. orthopsilosis* | Co1 | 9 | N/A | PRJNA322245 |
| 599 | Baltimore, USA | *C. orthopsilosis* | Co2 | 20 | N/A | PRJNA322245 |
| MSK616 | New York | *C. orthopsilosis* | - | 8[3] | N/A | This study |

| | | | | | | |
|---|---|---|---|---|---|---|
| MSK636 | New York | *C. orthopsilosis* | Co1 | 8 | N/A | PRJNA579121 |
| MSK638 | New York | *C. orthopsilosis* | Co1 | 8 | N/A | PRJNA579121 |
| MSK639 | New York | *C. orthopsilosis* | Co1 | 9 | N/A | PRJNA579121 |
| 748 | Atlanta, USA | *C. orthopsilosis* | - | 2 | N/A | PRJNA322245 |
| MSK805 | New York | *C. orthopsilosis* | - | 2 | N/A | This study |
| 831 | Baltimore, USA | *C. orthopsilosis* | Co1 | 9 | N/A | PRJNA322245 |
| 90-125 | San Francisco, USA | *C. orthopsilosis* | - | 2 | N/A | PRJNA431439 |
| B-8274 | Pakistan | *C. orthopsilosis* | - | 2 | N/A | PRJNA579121 |
| B-8323 | Pakistan | *C. orthopsilosis* | - | 2 | N/A | PRJNA579121 |
| MCO456 | San Antonio, USA | *C. orthopsilosis* | Co4 | 1 | N/A | PRJEB4430 |
| Engineered/evolved strains | | | | | | |
| Name | Parent | Description | | | | |
| 795B, B16 | MSK795 | Miltefosine-resistant evolved strains | | | | |
| 247A1,B1,C1,D1,D2, D16,E1, E16 | MSK247 | Miltefosine-resistant evolved strains | | | | |
| RTA3*Δ*/*Δ* 1 | CLIB214 | *rta3*Δ/*rta3*Δ (*cpar2_104610Δ* / *cpar2_104610*Δ) | | | | |
| RTA3*Δ*/*Δ* 2 | CLIB214 | *rta3*Δ/*rta3*Δ (*cpar2_104610Δ* / *cpar2_104610*Δ) | | | | |
| CPAR2_102700*Δ*/*Δ* 1 | CLIB214 | *cpar2_102700Δ* / *cpar2_102700Δ* | | | | |
| CPAR2_303950*Δ*/*Δ* 1 | CLIB214 | *cpar2_303950Δ* / *cpar2_303950Δ* | | | | |
| CPAR2_303950*Δ*/*Δ*, | CLIB214 | *cpar2_102700Δ* / *cpar2_102700Δ, cpar2_303950Δ* / *cpar2_303950Δ* | | | | |

[1]Newly reported strains isolated from the same patient (where known) are indicated using random patient numbers

[2]Copy number for strains with CNV-B was measured as number of copies of full ABC repeat (Supp. Fig 1)

[3]Sequence coverage data for this strain was too poor quality to call a CNV

Supplementary Table S2. List of Primers

| Primer: | Purpose: | Sequence: |
|---|---|---|
| **Deletion of RTA3 (CPAR2_104610)** | | |
| RTA3_KO_Guide_TOP | Oligo designed for cloning into pCP-tRNA vector as guide RNA sequence to cleave within RTA3 | CCACTGCTGTTGGAAGCCCATAT |
| RTA3_KO_Guide_BOT | Oligo designed for cloning into pCP-tRNA vector as guide RNA sequence to cleave within RTA3 | AACATATGGGCTTCCAACAGCAG |
| RTA3_KO_Upstream_Fwd | Primer designed for amplification of ~500b upstream of RTA3 | CAGAGGGTATCATTGGTG |
| RTA3_KO_Upstream_Rev | Primer designed for amplification of ~500b upstream of RTA3, includes inserted barcode for use as overlap in fusion PCR and screening transformants via colony PCR | CCCTGAAATGAGTGGTCTCTACTGGTCCATACTACTAG |
| RTA3_KO_Downstream_Fwd | Primer designed for amplification of ~500b downstream of RTA3, includes inserted barcode for use as overlap in fusion PCR and screening transformants via colony PCR | AGAGACCACTCATTTCAGGGCATAGGTAAATCTTTGGG |
| RTA3_KO_Downstream_Rev | Primer designed for amplification of ~500b downstream of RTA3 | TAGACCACACATGTTGCG |
| RTA3_Fusion_PCR_Fwd | Primer designed for use in fusion PCR of upstream and downstream amplicons | GGAAGTATCAATGTAAGCTG |
| RTA3_Fusion_PCR_Rev | Primer designed for use in fusion PCR of upstream and downstream amplicons | CATCAGTGGTTAGAGTCG |
| RTA3_Barcode_Fwd | Primer designed for amplifcation of product specific to RTA3 knockout barcode, for use with RTA3_KO_Downstream_Rev. For use in colony PCR | TGGACCAGTTAGAGACCAC |
| **Deletion of DNF1 (CPAR2_303950)** | | |
| 303950_KO_Guide_TOP | Oligo designed for cloning into pCP-tRNA vector as guide RNA sequence to cleave within CPAR2_303950 | CCAAAACGACACCTGCTGGGTGC |
| 303950_KO_Guide_BOT | Oligo designed for cloning into pCP-tRNA vector as guide RNA sequence to cleave within CPAR2_303950 | AACGCACCCAGCAGGTGTCGTTT |
| 303950_KO_Upstream_Fwd | Primer designed for amplification of ~700b upstream of CPAR2_303950 | CCTCATTCACTCCTTCTCTG |
| 303950_KO_Upstream_Rev | Primer designed for amplification of ~700b upstream of CPAR2_303950, includes inserted barcode for use as overlap in fusion PCR and screening transformants via colony PCR | TATGCAAATTCGTGCGTGTGCTTGAACAAACTAGGGTCTG |
| 303950_KO_Downstream_Fwd | Primer desgned for amplification of ~600b downstream of CPAR2_303950, includes inserted barcode for use as overlap in fusion PCR and screening transformants via colony PCR | CACACGCACGAATTTGCATAGTTGCATCGTATCAATGAAGG |
| 303950_KO_Downstream_Rev | Primer designed for amplification of ~600b downstream of CPAR2_303950 | TCCTTCGACGCCTGTTACTG |

| | | |
|---|---|---|
| 303950_Fusion_PCR_Fwd | Primer designed for use in fusion PCR of upstream and downstream amplicons | TCATTCACATACACGCACAC |
| 303950_Fusion_PCR_Rev | Primer designed for use in fusion PCR of upstream and downstream amplicons | GCTTTGTCTTTCCACAGTCC |
| 303950_Barcode_Fwd | Primer designed for amplifcation of product specific to CPAR2_303950 knockout barcode, for use with 303950_Fusion_PCR_Rev. For use in colony PCR | GTTCAAGCACACGCACGA |
| **Deletion of DNF2 (CPAR2_102700)** | | |
| 102700_KO_Guide_TOP | Oligo designed for cloning into pCP-tRNA vector as guide RNA sequence to cleave within CPAR2_102700 | CCAAATGGAACTGCAGCCAACCC |
| 102700_KO_Guide_BOT | Oligo designed for cloning into pCP-tRNA vector as guide RNA sequence to cleave within CPAR2_102700 | AAC GGG TTG GCT GCA GTT CCA TT |
| 102700_KO_Upstream_Fwd | Primer designed for amplification of ~500b upstream of CPAR2_102700 | CGCCTACTGGATCTGATTGA |
| 102700_KO_Upstream_Rev | Primer designed for amplification of ~500b upstream of CPAR2_102700, includes inserted barcode for use as overlap in fusion PCR and screening transformants via colony PCR | CTAAACAATGTCCGATCCGTGACAAACACACACACACTAC |
| 102700_KO_Downstream_Fwd | Primer designed for amplification of ~500b downstream of CPAR2_102700, includes inserted barcode for use as overlap in fusion PCR and screening transformants via colony PCR | ACGGATCGGACATTGTTTAGCCCAGCTTCTTGATGTTGAT |
| 102700_KO_Downstream_Rev | Primer designed for amplification of ~500b downstream of CPAR2_102700 | CCTCTTTCAACCACTACACC |
| 102700_Fusion_PCR_Fwd | Primer designed for use in fusion PCR of upstream and downstream amplicons | CAACAAAGGTCTCTTCTACCG |
| 102700_Fusion_PCR_Rev | Primer designed for use in fusion PCR of upstream and downstream amplicons | AGCATCAAAGCTTGGGTCA |
| 102700_Barcode_Rev | Primer designed for amplification of product specific to CPAR2_102700 knockout barcode, for use with 102700_Fusion_PCR_Fwd. For use in colony PCR | GAAGCTGGGCTAAACAATGTC |
| **RT qPCR of RTA3 and ACT1** | | |
| ACT1_Fwd | Primer used for amplification of ACT1 in qPCR | TGATGACGCACCAAGAGC |
| ACT1_Rev | Primer used for amplification of ACT1 in qPCR | GACCCATACCAACCATGATACC |
| RTA3_Fwd | Primer used for amplification of RTA3 in qPCR | CCTGTAGATGACGGGTATGAC |
| RTA3_Rev | Primer used for amplification of RTA3 in qPCR | TGCATCCCCAAGATCAGC |
